# First evidence of a causal link between genetic variation and thermal adaptation in a schistosome host snail

**DOI:** 10.1101/2024.01.02.573866

**Authors:** Tim Maes, Julie Verheyen, Bruno Senghor, Aspire Mudavanhu, Ruben Schols, Bart Hellemans, Enora Geslain, Filip A.M. Volckaert, Hugo F. Gante, Tine Huyse

## Abstract

Freshwater snails are pivotal in transmitting schistosomiasis, a tropical parasitic disease affecting over 150 million people. The adaptive potential of these snails is a critical factor in determining how climate change and other environmental factors influence disease transmission dynamics, yet it has remained unexplored. *Bulinus truncatus* is the schistosome intermediate host snail with the widest geographic distribution and therefore plays a pivotal role in determining the maximum range of urogenital schistosomiasis. In this study, we assessed the local adaptation capacity of *B. truncatus* to temperature through an integrative approach encompassing phenotypic, ecophysiological, and genomic data. Ten snail populations from diverse thermal environments were collected in three countries, with eight populations reared in a common garden. The F2 generation (total N= 2592) was exposed to eight chronic temperature treatments and various life-history traits were recorded for over 14 weeks. Subsequently, ecophysiological analyses were conducted on the ten last surviving snails per population. Genotyping the parental generation collected in the field using a genotyping-by-sequencing (GBS) approach, revealed 12,875 single nucleotide polymorphisms (SNPs), of which 4.91 % were potentially under selection. We observed a significant association between these outlier SNPs, temperature, and precipitation. Thermal adaptations in life-history traits were evident, with lower survival rates at high temperatures of warm- origin snails compensated for by higher reproduction rates. Cold-origin snails, on the other hand, exhibited higher growth rates adapted to a shorter growing season. Ecophysiological adaptations included elevated sugar and haemoglobin contents in cold-adapted snails. In contrast, warm-adapted snails displayed increased protein levels but also more oxidative damage. Furthermore, heightened phenoloxidase levels indicated a more robust immune response in snails from parasite-rich regions. The substantial local adaptation capacity of *B. truncatus* holds profound implications for its response to climate change, future schistosomiasis risk, and the effectiveness of schistosomiasis control measures.

**Highlights:**

- Local adaptation influences species’ responses to climate change
- The snail *Bulinus truncatus* showed a high thermal local adaptation (LA) potential
- LA is apparent through variations in life history and ecophysiological traits
- We identified a significant genetic basis underlying this LA
- LA of the hosts could sustain schistosomiasis transmission under global warming

## 1 Introduction

Climate change, characterised by heightened temperatures and shifts in precipitation patterns, profoundly impacts freshwater ecosystems by modifying key waterbody characteristics (IPCC, 2022; Knouft & Ficklin, 2017). These changes have a large effect on freshwater organisms, including disease vectors like mosquitoes and freshwater snails. Disease vectors faced with climate change must either adapt to their changing surroundings or shift their distribution to align with current environmental conditions (Berg et al., 2010; Jump & Peñuelas, 2005; Williams et al., 2008), thereby significantly impacting the global dynamics of vector-borne diseases (Ryan et al., 2020; Stensgaard et al., 2019; Williams et al., 2016).

The capacity of organisms to adapt to climatic conditions increases the fitness of populations in their native environments (Blanquart et al., 2013; Savolainen et al., 2013), but it can also pose challenges to the long-term survival of populations under climate change (DeMarche et al., 2019). Local adaptation is evident through inter-population variations in life history traits, such as body size (Daufresne et al., 2009; James, 1970), growth rate (Olsson & Uller, 2003) or survival (Seefeldt & Ebert, 2019). Additionally, these adaptations may manifest more subtly in ecophysiological traits (Verheyen & Stoks, 2019). Strong local adaptation in widely distributed species may result in a narrower environmental tolerance of the local populations compared to the species as a whole (Holt, 2009). Consequently, these locally adapted populations could be more vulnerable to climate change if the rate of change surpasses the species’ dispersal and adaptation capacity (DeMarche et al., 2019). Therefore, assessing the effects of climate change on disease vector distributions while assuming populations lack local adaptation may lead to inaccurate estimates of range changes (Atkins & Travis, 2010; DeMarche et al., 2019; Valladares et al., 2014). Integrating species life history data, landscape genetic data and species-specific ecophysiological data into species distribution models has the potential to significantly refine vector distribution predictions (Aleuy et al., 2023; DeMarche et al., 2019; Razgour, 2015), thereby improving estimates of disease risk.

Freshwater snails play a pivotal role in the transmission of schistosomiasis, a tropical disease affecting over 150 million people worldwide, the majority living in Sub-Saharan Africa (WHO, 2015). Besides the significant health threat, the disease has a large economic burden on communities as it is both a cause and a consequence of poverty (Rinaldo et al., 2021). Among the transmitting snails, *Bulinus truncatus* (Audouin, 1827) stands out as a key species transmitting both human urogenital and bovine schistosomiasis. It is believed to have the broadest distribution range among all intermediate host snails, spanning across the entire African continent, Southern Europe and the Middle East (Brown, 1994), thus determining the maximal geographic spread of the disease.

Although local adaptation to climate is common in species with wide distribution ranges (Bocedi et al., 2013; Jump & Peñuelas, 2005; King et al., 2018), studies on schistosomiasis host snails are limited and often characterised by flawed experimental designs, small sample sizes and pseudo-replication (Maes et al., 2021). Nonetheless, *B. truncatus* displays some spatial variation in life history traits (Diakité et al., 2023; Konan et al., 2022; Mulero et al., 2019), and molecular research on this species indicates significant neutral genetic differentiation and genetic diversity that is structured via isolation by distance on both intercontinental and regional scales (Chlyeh et al., 2002; Maes et al., 2022; Zein-Eddine et al., 2017). These findings collectively suggest a high potential for local adaptation in this snail (Chung et al., 2023; Savolainen et al., 2013) that might influence this species’ response to climate change, and consequently, schistosomiasis transmission dynamics.

In this respect, this study investigates the local adaptation potential of *B. truncatus* to temperature through an integrative approach including (i) a common garden experiment with different chronic temperature treatments to assess life history and ecophysiological trait variations among populations, and (ii) a landscape genomics approach using genotyping by sequencing (GBS) to associate single-nucleotide polymorphisms (SNPs) with climatic variables and the obtained life history data. While the common garden experiment should control for phenotypic plasticity (Valladares et al., 2014; Yampolsky et al., 2013), and the use of second-generation offspring should limit environmental maternal effects, potential plastic responses induced by the common garden environment may bias inferences on adaptive divergence (Gienapp et al., 2008). Therefore, we also conducted a genotype-environment analysis to screen the genome for signs of adaptive differentiation, unconfounded by phenotypic plasticity. Moreover, given the polygenic control of most phenotypic traits, with many loci exerting small effects (Barghi et al., 2019; Savolainen et al., 2013), population genetic screens may overlook signatures of adaptive differentiation in such traits (Hoban et al., 2016; King et al., 2018). Therefore, the complementary approach used in this study, integrating genomic, phenotypic and landscape information, yields more reliable estimates of the local adaptation capacity of *B. truncatus*.

## 2 Materials and methods

### 2.1 Study species

The freshwater snail *B. truncatus* (Gastropoda, Heterobranchia) is a eurotypic species with very wide tolerance limits to abiotic factors, self-fertilisation capacities and a high reproduction rate (Appleton, 1978). It can easily be dispersed on the feet and feathers of birds (Pfenninger et al., 2011) or through passive dispersal in water currents or anthropogenic vectors (Kappes & Haase, 2012), thereby overcoming dispersal barriers. The snail is a tetraploid (2n=72) species that presumptively originated through genome duplication (autoploidy) with limited genomic divergence between the two genomes (Young et al., 2022). Although polyploidy can have detrimental effects on fertility and fitness (Van De Peer et al., 2017), it has also been linked to facilitating adaptation and ecological resilience, enabling polyploids to colonize new or rapidly changing environments (David, 2022). The multiple gene copies in the tetraploid genome could increase genetic diversity (Heslop-Harrison et al., 2023), and create the opportunity for adaptation (Comai, 2005).

### 2.2 Field collection and experimental design

*Bulinus truncatus* snails were collected in the second half of 2021 from ten locations in three countries spanning a latitudinal and temperature gradient (Table 1, Fig 1). Additionally, a French (Corsican) strain that has been maintained in the lab since 2014 was included. We chose French sampling sites on the border between the Corsican montane broadleaf and mixed forests and the Tyrrhenian-Adriatic Sclerophyllous and mixed forests ecoregions in the hot summer Mediterranean climate as this area represents the cold limit of the distribution range of *B. truncatus*. This area is characterised by a high seasonality with average air temperatures ranging between 7 °C in winter and 27 °C in summer. Senegalese snails were collected in the Sahelian Acacia savanna ecoregion characterised by a hot desert climate which represents the warm limit of the distribution range with less strong seasonality and average air temperatures ranging from 30 to 39 °C. Finally, the snails from Zimbabwe originate from the Zambezian and Mopane woodlands (Triangle and Malilangwe populations, characterised by a hot, semi-arid steppe climate) and the Southern Miombo woodlands (Imire, characterised by a subtropical highland climate) ecoregions where temperatures show less seasonality and are more temperate with averages between 14 and 22.6 °C. The field-collected snails were bred until the second (F2) generation in a common garden at 24 °C and a 12:12 h day-night regime, fed *ad libitum* with pesticide-free lettuce (dried at 55 °C for 12 h), and green macro-algae were added to enrich the water with oxygen. An overview of the different experiments and the endpoints measured is given in Fig 2.

**Table 1.**
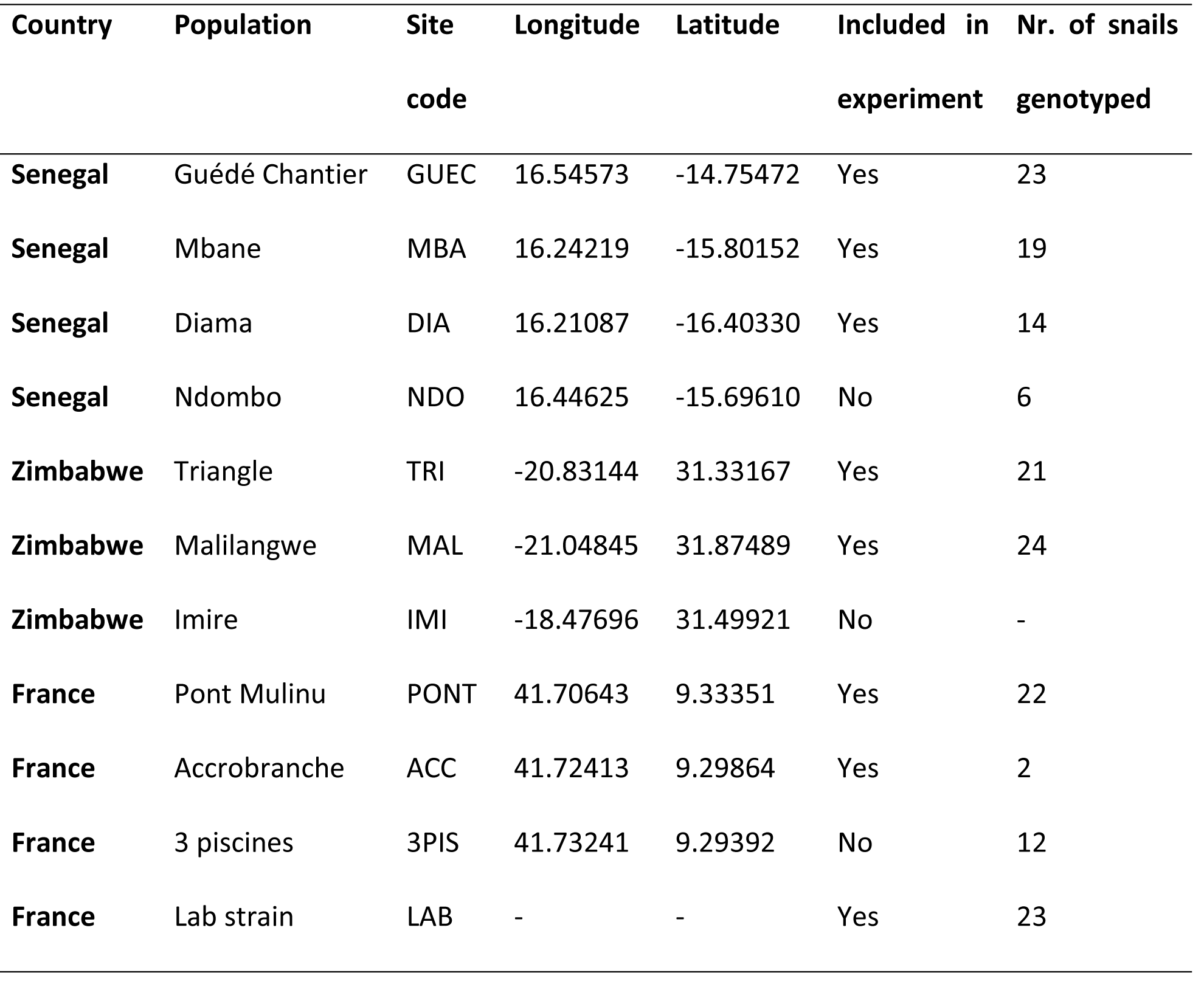
**The origins of the experimental snails**, together with the site codes, the coordinates of the respective locations, whether the population was included in the experiment and the number of snails successfully genotyped.

**Figure 1.**
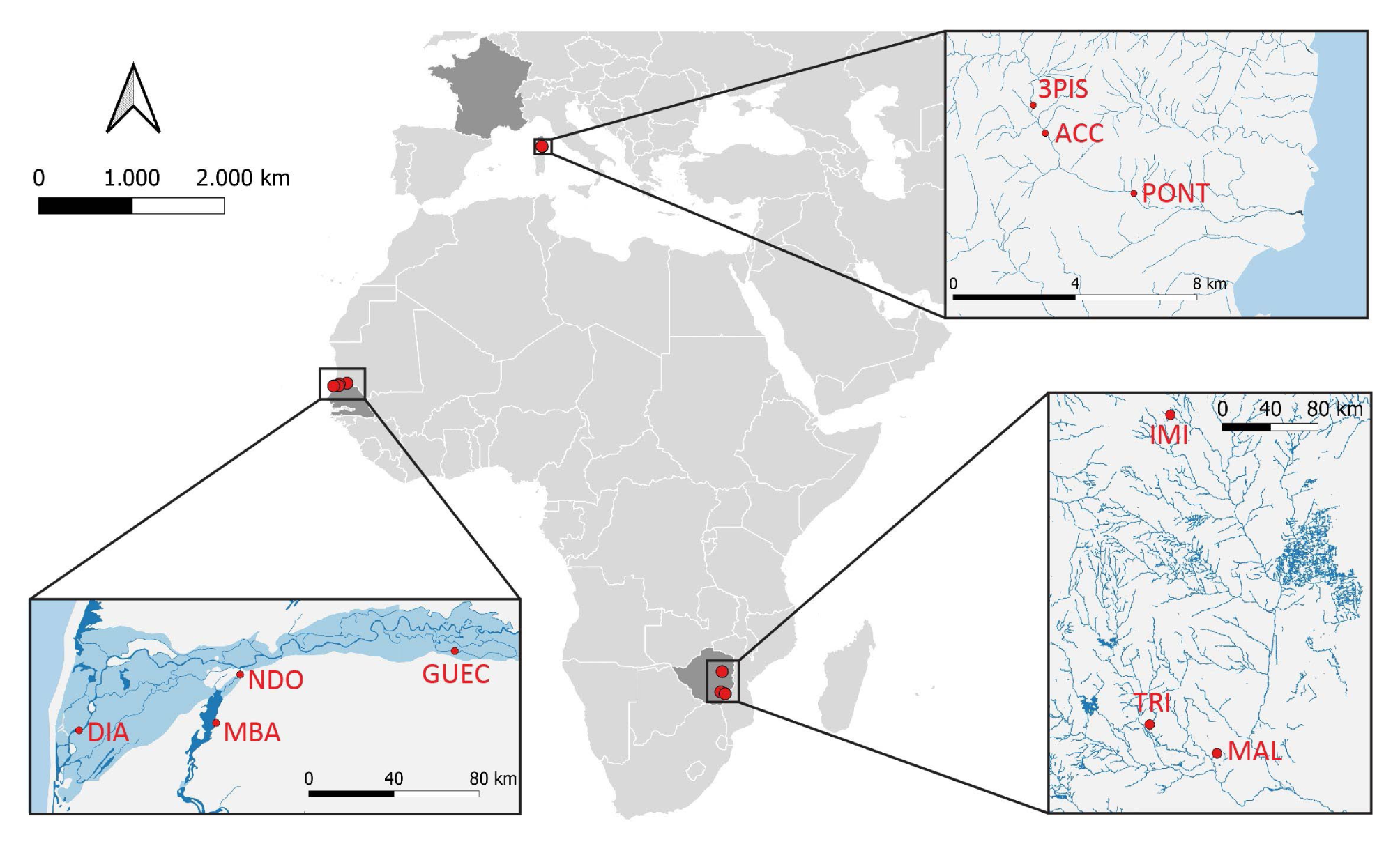
Locations of the sampling sites. The full names of the populations are given in Table 1.

**Figure 2.**
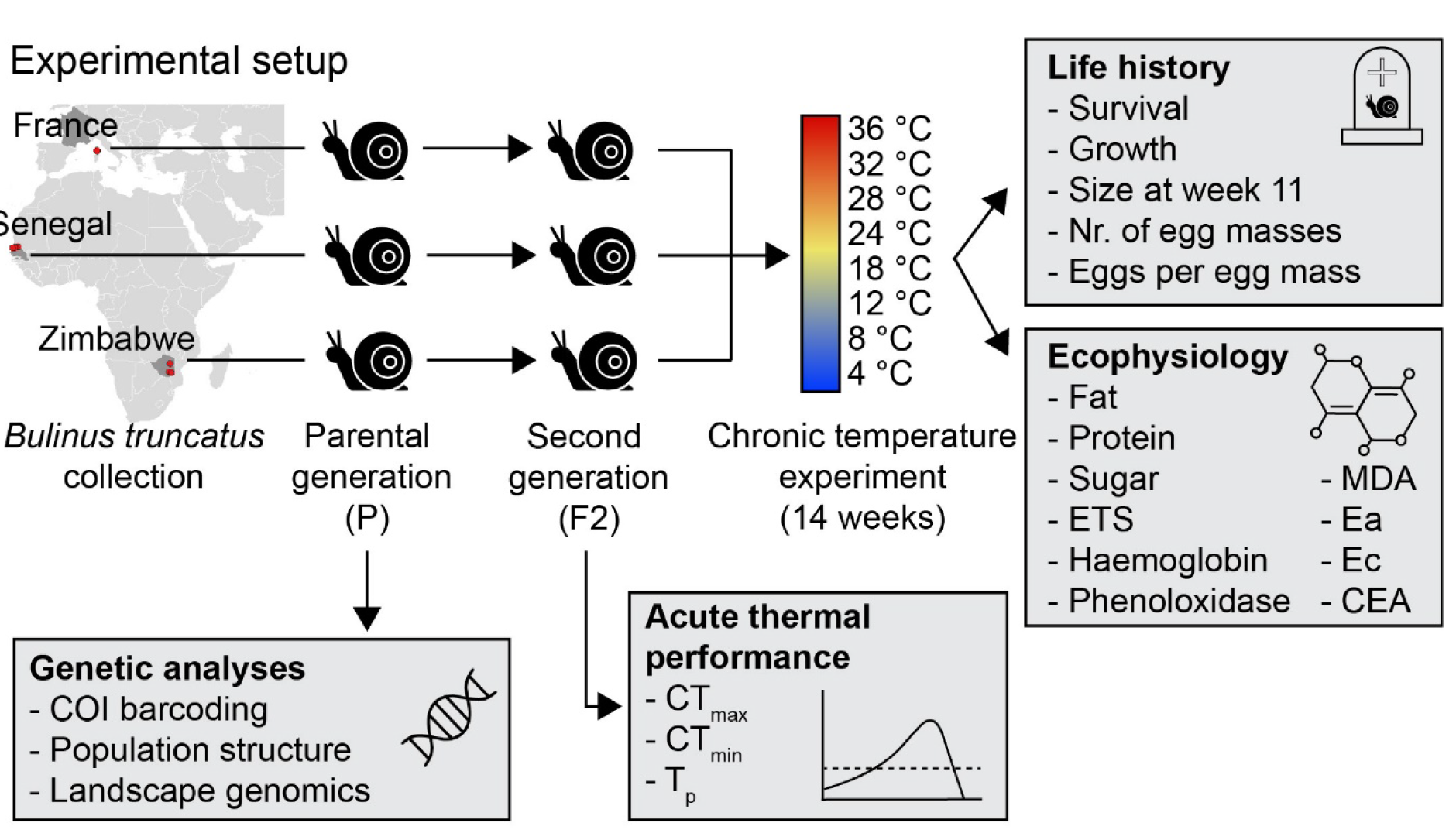
Overview of the experimental setup. Snails were caught in three different ecoregions (Fig 1) and bred in the lab until the second (F2) generation (the first F1 generation is not shown in this figure). Subsequently, the parental (P) generation snails caught in the field were analysed genetically. The second (F2) generation offspring were subjected to eight different chronic temperatures for 14 weeks. During this time various life history parameters were measured weekly (except for size at week 11) and the ten last surviving snails per location and treatment combination were analysed ecophysiologically. A subset of F2 snails that were not included in the chronic temperature experiment were used to asses the acute thermal performance of the snails.

DNA was extracted from the whole body of the parental (P) generation collected in the field using the E.Z.N.A.® mollusc DNA kit according to the manufacturer’s protocol (Omega Bio- tek). To verify the morphological identification from the field, a subsample of five snails from each population was genotyped through DNA barcoding using the protocol described in Maes et al. (2022) and identification of the specimens relied on a BLAST search against the NCBI database GenBank. After DNA barcoding, eight populations from three different countries identified as *B. truncatus* (see results section) were selected for the life history experiment (Table 1) while all populations identified as *B. truncatus* were included in the genetic analyses. The Imire population from Zimbabwe was identified as *Bulinus tropicus* (Krauss, 1848) and excluded from the experiment and genetic analyses.

The F2 snails were individually placed in 100 ml plastic cups filled with aged tap water at four to eight weeks old. The experiment was started in 12 different batches, each batch containing three snails per location for each temperature treatment (3 snails x 9 locations x 8 temperatures x 12 batches = 2592 snails in total, 36 snails/population/temperature). The temperature was increased/decreased by 2 °C/day starting from 24 °C until the snails reached their experimental temperature (4, 8, 12, 18, 24, 28, 32 or 36 °C) at the start of the experiment (time t=0). A constant temperature was permanently monitored using Hobo onset data loggers (tidbit v2 Temp logger). The snails were fed *ad libitum* with dried lettuce and green algae, and the water was refreshed weekly or when the oxygen level dropped below 5 mg/ml. The lettuce was changed daily in the high-temperature treatments (24, 28, 32 and 36 °C) to prevent its decomposition and associated oxygen depletion. The snails were kept at the experimental temperatures for 14 weeks.

### 2.3 Response variables

Each week the number of egg masses per snail was counted, dead snails were counted and removed, and shell height (apex to bottom of aperture) was measured under a stereomicroscope (Ceti Steddy-B) with a built-in ruler. The ten last surviving snails per treatment and location combination were taken out of their shell, the body was weighed with an accuracy of 0.01 mg (Mettler Toledo AB135-S, Columbus, Ohio, USA) and stored at -80 °C for ecophysiological analyses. Egg masses were collected in weeks one to seven and in week 12 and stored in 98 % ethanol to later quantify the number of eggs per clutch under a stereomicroscope.

The critical thermal maximum and minimum (CTmax and CTmin, respectively; Bartnicki et al., 2021; Morgan et al., 2018) were measured on a separate subset of F2 snails that were not included in the chronic temperature experiment. For measuring the CTmax, the snails were individually put in 12 ml plastic tubes filled with aged tap water and placed in a heating block (Thermo Fisher Scientific). The water temperature increased by 0.1 °C/min (following Johansson & Laurila, 2017). For measuring the CTmin, the snails were also individually put in 12 ml plastic tubes filled with aged tap water and placed in a water bath. The water temperature was decreased by 0.1 °C/min using a cooler. Before both runs, the size of each snail was measured under a stereomicroscope with a built-in ruler. Snails were considered fainted when they showed no response to tactile stimuli (i.e. the snails did neither retreat in their shell nor retreat their tentacles). After the test, snails were allowed to recover and only snails that fully recovered within 10 min. were considered for the analysis (5 out of 120 snails died in the CTmax experiment, none in the CTmin experiment).

To create a temperature gradient to assess the preferred temperature (Tp), ten aluminium U- profiles (lanes) of two meters in length were attached to create ten replicas and filled with aged tap water (cf. Johansson & Laurila, 2017). One side of the profiles was placed on a cooling plate, and the other side on a heating plate. The temperatures of both plates were adjusted to generate a stable temperature gradient of 24 °C in the middle of each lane and an increase/decrease of 1 °C/10 cm in the direction of the heater/cooler (min: 14 °C max: 34 °C). Ten snails that were not included in the chronic temperature experiment were randomly selected from each location and randomly assigned to one of the ten lanes. The snails were placed in the middle at 24 °C and the temperature at the position of each snail was measured every 15 min for 4 h. The first 2 h were considered as acclimation time while the preferred temperature was defined as the average temperature during the last two hours. We assumed that there were no differences in movement speed between the populations (Dillon et al., 2012).

### 2.4 Ecophysiological analyses

Whole-body snail samples stored at -80 °C were homogenised, 15 X diluted in phosphate-buffered saline (PBS) and centrifuged at 4 °C for 10 min. at 13,000 g. A minimum of 8 mg snail wet weight was required to run all ecophysiological analyses. If this weight was not reached, we pooled two snails from the same location and temperature treatment to obtain the minimum required weight.

Various ecophysiological parameters were examined to gain a comprehensive understanding of the snail’s overall well-being and to identify nuanced signs of local adaptation that might not be evident through life history traits alone. Assessing the snails’ general condition involved determining fat, total sugar, and protein contents (for the full protocols, see Appendix 1). From these variables, the cellular energy allocation (CEA) was calculated as outlined in Gomes et al. (2015). The CEA integrates the energy available (Ea) and energy consumption (Ec) of an organism. Additionally, the haemoglobin content was measured to gauge the oxygen-binding capacity. It should be noted that haemoglobin is absent in most freshwater snails, except for pulmonate species like *B. truncatus* (Lieb et al., 2006). Furthermore, the malondialdehyde (MDA) levels were quantified to assess oxidative damage to lipids (Miyamoto et al., 2012). Phenoloxidase (PO), instrumental in the immune response to various parasite species in snails (Le Clec’h et al., 2016), was also measured to evaluate the strength of the snails’ immune system. Lastly, the snails’ metabolic rate was quantified at the cellular level by assessing the activity of the electron transport system (ETS) (De Coen & Janssen, 2003). Detailed protocols for quantifying these variables are provided in Appendix 1.

Sample sizes for all variables analysed ranged from one (for some localities in the 32 °C treatment) to 22 with an average of 8.7 samples per population and temperature treatment (Table A1).

### 2.5 Data analyses

All statistical analyses were performed in R 4.2.2 (R Core Team, 2022). Growth rates were calculated for different time points (weeks 0-3, 3-6, 6-9 and 9-11) and the effect of country of origin and temperature on growth rates and the shell size at week 11 were assessed using a linear mixed effects model (‘lmer’ package). For the effects of the explanatory variables on the number of egg masses laid, a generalised linear mixed effects model was built with a zero-inflated Poisson distribution and a logit link function (R package ‘glmmTMB’). The initial size of the snails was added as a covariate and population nested in country, individual snails nested in population (since egg masses have been collected at multiple time points for some snails), and batch number were added as random factors in the egg masses, growth rate, and snail size models. Only data from the first 11 weeks were used for all analyses as from week 12 onwards all Senegalese snails had died at 28 °C and no data were available for weeks 12 to 14 in this treatment. To assess the effects of country of origin and temperature on the number of eggs per egg mass, a generalised linear mixed effects model with a Poisson distribution was built. The snail size at the time of egg laying was added as a covariate, and population nested in country and batch number were added as random factors. The effects of country and temperature on the survival of snails were tested using Cox proportional hazards regression (R package ‘survival’). Since it is not possible to add random factors to a Cox proportional hazards regression, both the initial snail size and batch number were included as covariates in this analysis. Population was also added as a covariate but this factor was not significant. The effects of temperature, country of origin and the number of weeks survived (only for fat) on the ecophysiological parameters were analysed using either linear mixed-effects models (fat and PO) or linear models (protein, sugar, ETS, MDA, haemoglobin and CEA; ‘lmer’ package).

Fat, MDA and haemoglobin contents were log-transformed while PO activity, sugar content, ETS activity and CEA were square root transformed to obtain a normal distribution of the model residuals. The goodness of fit of the models was assessed using the Akaike Information Criterion (AIC). Assumptions of homogeneity of variance of the residuals and normality of the residuals were met for all linear (mixed-effects) models (Zuur, Ieno, & Elphick, 2010). No overdispersion or multicollinearity among the explanatory variables was observed in the zero-inflated Poisson model while the Cox-proportional hazards model was tested for the proportional hazards assumption, influential outliers and nonlinearity in the relationship between the log hazard and the covariates (R package ‘survminer’). Wald χ^2^, *F*-statistics, and accompanying *p*-values of the fixed effects were calculated, and significant interactions were further evaluated using pairwise contrasts of estimated marginal means through the ‘emmeans’ package for all analyses.

The effect of country of origin on CTmax, CTmin and Tp values was assessed using an analysis of covariance (ANCOVA) with snail size as a continuous covariate and country of origin as the categorical variable. To meet the assumption of normality, the CTmin data was log-transformed. Pairwise comparisons between countries were carried out using Tukey’s post hoc comparison tests.

### 2.6 SNP genotyping

DNA from the parental (P) generation snails was used to build a paired-end GBS reduced representation library (RRL) according to Elshire et al. (2011) and sequenced on an Illumina NovaSeq 6000 platform (Genomics Core KU Leuven). High molecular weight DNA was digested with NsiI as the restriction enzyme and Illumina sequencing primers P1 and P2 and adapters containing the barcodes were ligated to the resulting fragments.

The NovaSeq run produced 2.01 x 10^9^ paired-end raw 101 and 92 bp reads (forward and reverse reads, resp.). Reads were demultiplexed using process_radtags from Stacks 2.5 (Catchen et al., 2013) and the sequence quality was checked using FastQC (http://www.bioinformatics.babraham.ac.uk/projects/fastqc). FastX trimmer (http://hannonlab.cshl.edu/fastx_toolkit/) was used to trim the first 7 bp of the forward reads and the first 6 bp of the reverse reads. Trimmed forward and reverse reads were mapped against the *B. truncatus* reference genome (Young et al., 2022) using Bowtie2 (Langmead et al., 2019) and SNPs were identified using GATK UnifiedGenotyper 3.7. The resulting vcf file containing all variants was further filtered to only include biallelic SNPs, a read depth between 20 and 100, a minimum genotyping quality of 20, a minimum allelic depth of 6, a maximum of 40 % missing data per locus, a maximum of 70 % missing data per individual and a minor allele frequency of 0.02. The number of successfully genotyped individuals is given in Table 1. Since *B. truncatus* is presumed to be allotetraploid (Young et al., 2022), a ploidy test was carried out using nQuire (Weib et al., 2018). NQuire uses next-generation sequencing data to distinguish between different ploidy levels based on the frequency distributions at variant sites where only two bases are segregating. SNP identification was carried out with both diploid (n=2) and tetraploid (n=4) settings in UnifiedGenotyper and a PCA was carried out on both filtered vcf files to check if the two datasets gave similar outcomes (Fig A1). Since both datasets gave the same output, all downstream analyses were carried out using the diploid dataset since more analysis tools are available for diploid data.

### 2.7 Outlier detection

Two methods were used to identify outlier SNPs putatively under natural selection; a SNP was considered putatively adaptive when it was identified as an outlier by both methods. Firstly, outliers were detected based on the Bayesian likelihood approach implemented in BAYESCAN 2.1 (Foll & Gaggiotti, 2008). BAYESCAN 2.1 identifies candidate loci under natural selection from genetic data, using differences in allele frequencies between populations, thereby reducing the number of false positives considerably (Narum & Hess, 2011). BAYESCAN 2.1 was run for 10,000 iterations and a burn-in of 200,000 steps. The prior odds of neutrality parameter (pr_odds) was set to 10,000 (Lotterhos & Whitlock, 2014) and the false discovery rate (q-value) to 0.01. Secondly, an individual-based method, Pcadapt v4 (Privé et al., 2020), was used that assumes that candidate markers are outliers with respect to how they are related to population structure as represented by a principal component analysis (PCA). For this, a PCA was performed and a screeplot in combination with a STRUCTURE analysis (see below) was used to choose the number of principal components (*K*=5) to retain. The package then regressed all variants onto the resulting principal components to get a matrix of Z-scores to integrate all PCA dimensions in one multivariate distance for each variant. These distances approximately follow a chi-squared distribution, which enabled the derivation of one *p*-value for each genetic variant. A Bonferroni correction was applied to identify outlier SNPs. An outlier dataset, containing all SNPs that were identified by both methods and a neutral dataset, without these outlier SNPs, were constructed to be used in subsequent analyses.

### 2.8 Population and landscape genomics analyses

The neutral dataset was used to assess pairwise *Fst* values between populations using 10,000 permutations. Genetic clustering among populations was assessed using a PCA on both the neutral and outlier datasets. Population structure was estimated using a Bayesian Markov Chain Monte Carlo (MCMC) model implemented in STRUCTURE v2.3.4 (Pritchard et al., 2000) using 10 replicates for each number of populations (*K*= 4-7) with a burn-in period of 50,000 and 100,000 repetitions. The most probable *K*-value was determined using Structure Harvester (Earl & VonHoldt, 2012) based on the Δ *K*-value (Evanno et al., 2005). Allelic richness, the observed (*Ho*) and expected heterozygosity (*He*), and inbreeding coefficients per population were calculated using the R package ‘hierfstat’ (Goudet, 2005). A latent factor mixed model (LFMM) was run using the R package ‘lfmm’ (Caye et al., 2019) on both the full and outlier SNP datasets to identify loci putatively subject to adaptive selection. LFMMs detect correlations between environmental and genetic variation while simultaneously inferring background levels of population structure (Frichot et al., 2013). The number of clusters (*K*=5) used as input for the LFMM was inferred from the PCA and STRUCTURE analyses (see below). Finally, a population-based redundancy analysis (RDA) was carried out using the R package ‘vegan’ (Oksanen et al., 2022) on the allele frequencies of the populations to detect loci putatively under selection (Forester et al., 2018). For each of the 19 available bioclimatic variables (extracted from worldclim.org), values were extracted for each population on a five km resolution. Environmental variables were scaled and centred; multicollinearity among variables was checked using the Pearson’s correlation coefficient and the variance inflation factor (VIF). Only variables with a correlation coefficient <0.70 and a VIF <10 were retained for the RDA and consisted of the annual temperature range (bio7), the mean temperature of the driest quarter (bio9), precipitation of the wettest month (bio13) and precipitation seasonality (bio15). A backward selection procedure was followed to determine relevant environmental variables and allele frequencies were Hellinger-transformed before the RDA. The adjusted *R*^2^ was calculated to correct for the number of explanatory variables and the significance was determined using 5,000 permutations. SNPs under selection were identified from the loadings of the significant RDA axes as those with values ±1 standard deviation (*p*=0.05) from the mean loading. This procedure was then repeated to detect loci explaining the variance in life history traits. Only traits that showed a significant difference between the countries of origin were included in the analysis. The means per population for each trait and temperature treatment, and the allele frequencies per population included in the life history experiment were used. Subsequently, the same procedure as for the environmental layers was used to carry out the RDA analysis.

## 3 Results

### 3.1 Molecular identification

All populations were identified as *B. truncatus* based on molecular barcoding, except for the Imire population (Zimbabwe) which was identified as *Bulinus tropicus*, and hence removed from all other analyses. Furthermore, the Zimbabwean population from Malilangwe clustered apart from all other populations in the PCA analysis (Fig 6). Because the Malilangwe population was identified as *B. truncatus* both through COI barcoding and shell morphology, and identified as tetraploid by the nQuire analysis (Appendix 2), this population was not excluded from this study. Instead, the Malilangwe population was treated as a separate origin and not taken together with the Triangle population.

### 3.2 Life history traits

The temperature had a significant effect on the survival of the snails (Fig 3, *W*_43_=1164, *p*<0.001). At 4 and 36 °C, all snails were dead after one week (Fig 3a,h) and at 32 °C after two weeks (Fig 3g). At 8 °C, mortality was very low for all countries (<20 % after 14 weeks, Fig 3b), except for the Malilangwe snails where mortality reached 68 % after 11 weeks (contrast: *p*<0.001). This population also had higher mortality rates than Zimbabwean (contrast: *p*=0.0376) and French snails (contrast: *p*=0.0388) at 12 °C (Fig 3c). At 24 °C, the mortality of the Senegalese snails was higher than snails from France (contrast: *p*=0.0074), Zimbabwe (contrast: *p*<0.001) and Malilangwe (contrast: *p*<0.001, Fig 3e). All countries had high mortality rates at 28 °C (>62 % after 14 weeks) with Zimbabwean snails having lower mortality rates than snails from France (contrast: *p*=0.0153), Malilangwe (contrast: *p*=0.0034) and Senegal (contrast: *p*=0.0017, Fig 3f).

**Figure 3.**
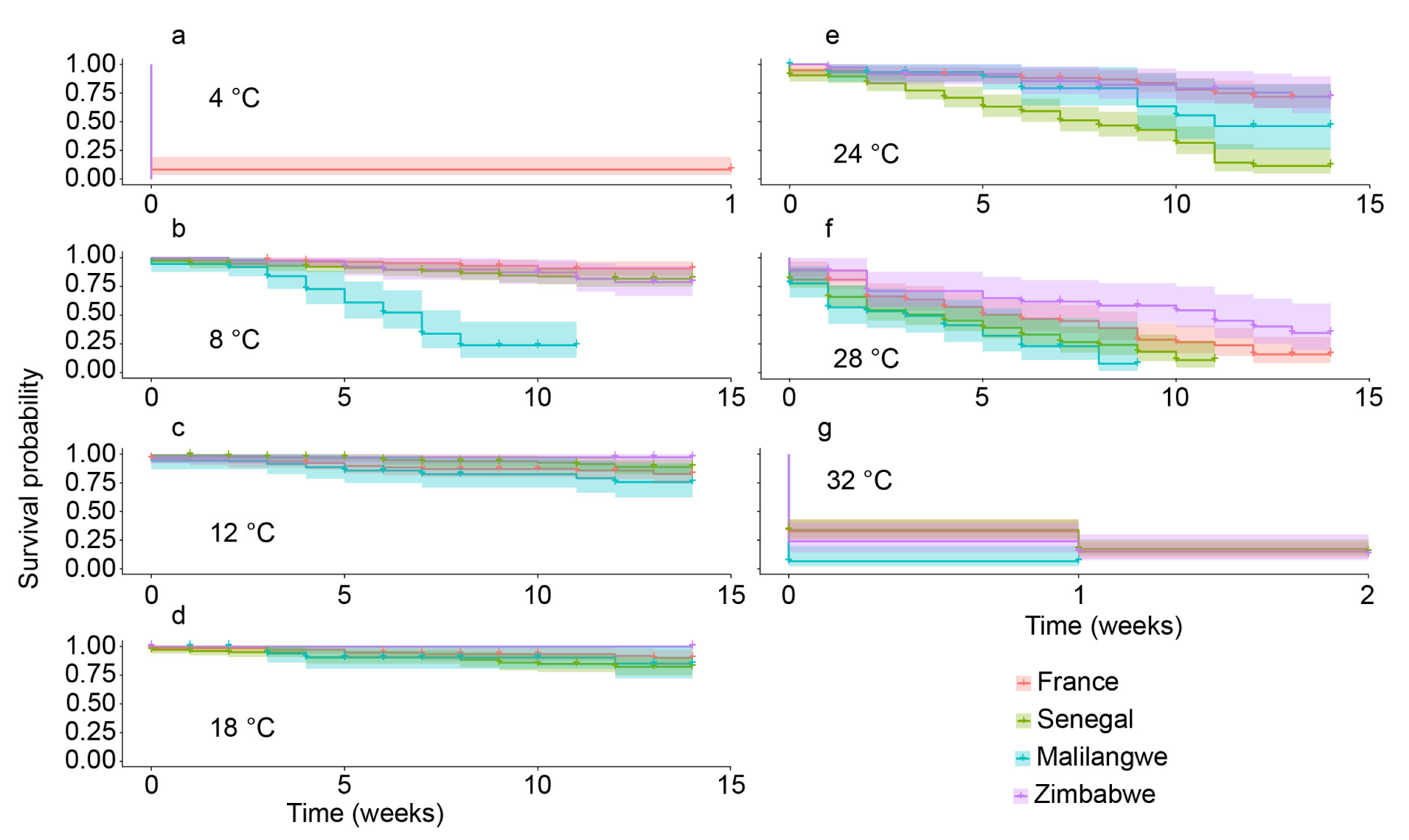
Survival curves as a function of country of origin and temperature (a-g).

Growth rates differed significantly between countries (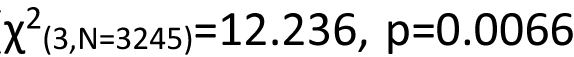, Fig 4a). Snails from Corsica had higher growth rates than Malilangwe (contrast: *p*=0.0112) and tended to have higher growth rates than Zimbabwe (contrast: *p*=0.0772, Fig 4a). Snail growth rates increased with temperature 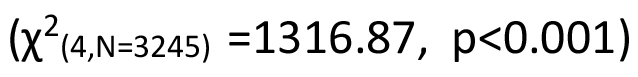 but no significant differences could be observed between the 18 and 24 °C treatments (contrast: *p*=0.3145). Finally, there was an interaction between the country of origin and temperature 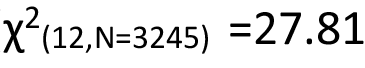, *p*=0.0059), with Malilangwe snails having lower growth rates at 8 °C (contrast: *p*=0.0139) and 18 °C (contrast: *p*=0.0472) than French snails, and Zimbabwean snails having lower growth rates at 28 °C than both French (contrast: *p*=0.0004) and Senegalese (contrast: *p*=0.0371) snails. There were no significant differences in growth rates between countries at other temperatures. Snail sizes at week 11 were significantly different between countries of origin (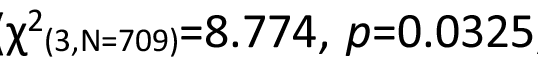, Fig 4b) although the *post-hoc* test lacked the power to pick up these differences (contrast France-Malilangwe *p*=0.0902).

**Figure 4.**
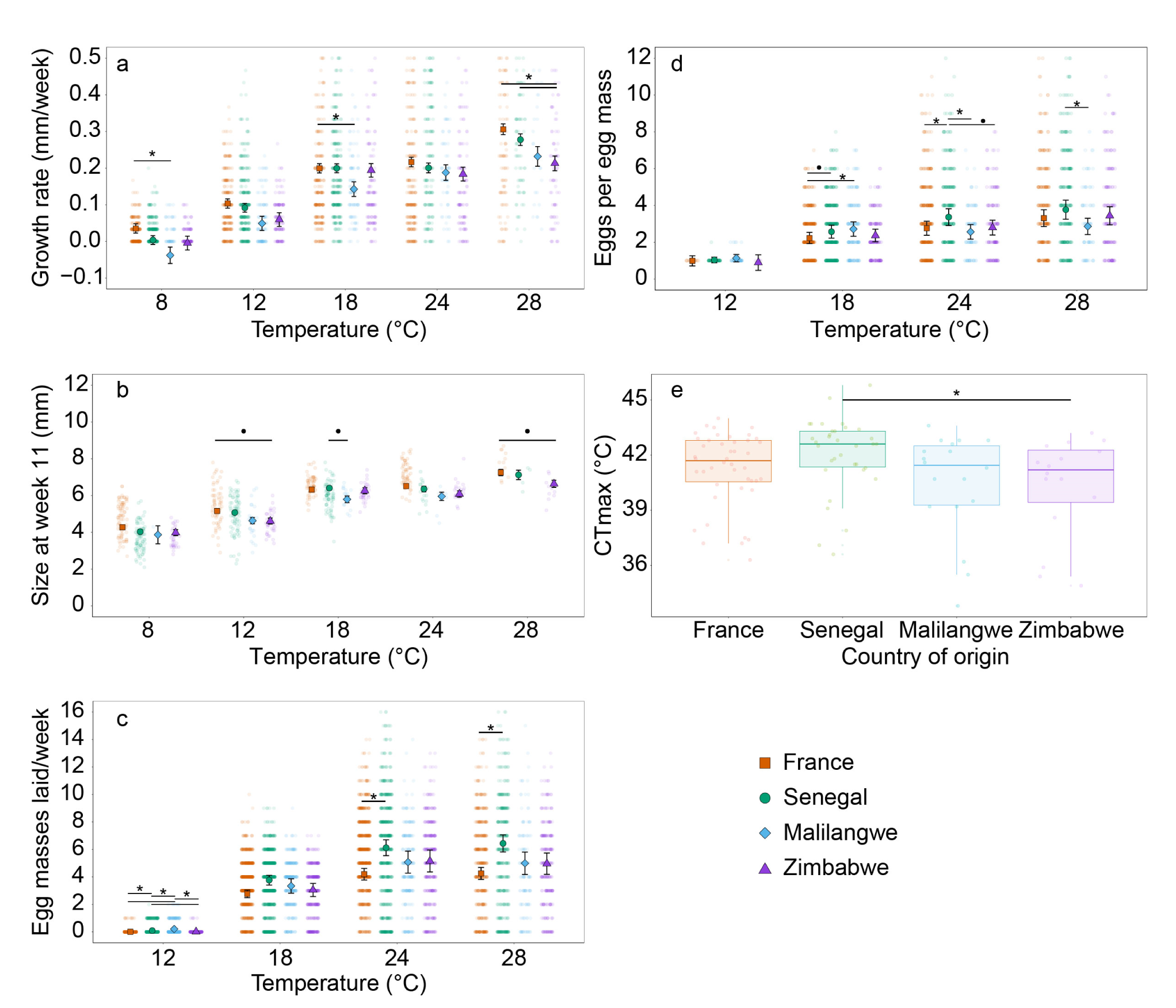
Results of the life history analyses. a) Growth rate, b) The snail size at week 11, c) The number of egg masses laid, and d) The number of eggs per egg mass as a function of temperature and country of origin, e) The CTmax, as a function of country of origin. Significant (p<0.05) main or interaction effects of temperature and country of origin are indicated by asterisks. Trends are indicated by bullets.

The country of origin influenced the number of egg masses laid (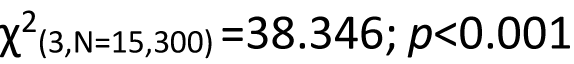, Fig 4c). Specifically, French and Zimbabwean snails laid fewer egg masses than Senegalese (contrasts: *p*<0.001 for France and *p*=0.0087 for Zimbabwe) and Malilangwe snails (contrasts: *p*<0.001 for France and *p*=0.0144 for Zimbabwe). Furthermore, temperature also impacted the number of egg masses laid 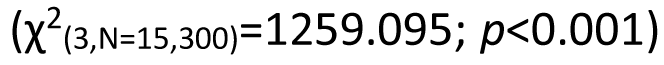. Firstly, no eggs were laid at temperatures below 8 °C or above 32 °C. Secondly, the number of egg masses was lower at 12 °C compared to 18, 24 and 28 °C (all contrasts: *p*<0.001). Thirdly, most egg masses were laid at 24 and 28 °C (no in-between differences: contrast: *p*=0.999) while slightly fewer eggs were laid at 18 °C (contrast: *p*<0.001). Additionally, the number of eggs per egg mass tended to be influenced by the interaction between the country of origin and temperature (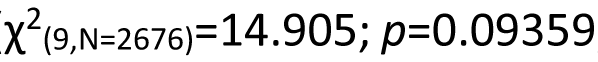, Fig 4d). The Senegalese snails tended to have a higher number of eggs per egg mass than French at 24 °C (contrast: *p*=0.0075), than Zimbabwean at 24 °C (contrast: *p*=0.0524), and than snails from Malilangwe at 24 °C (contrast: *p*=0.0131) and 28 °C (contrast: *p*=0.0126). The number of eggs per egg mass increased with temperature for all countries (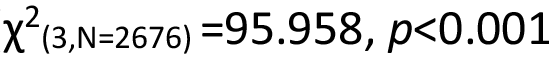, Fig 4d).

Finally, the country of origin had a significant influence on the CTmax of the snails (F3,107=2.898, *p*=0.0385). Snails from Senegal had a slightly higher CTmax than snails from Zimbabwe (contrast: *p*=0.047, Fig 4e), while there were no significant differences among the other countries. No significant differences between countries were found in CTmin (*F*3,116=2.250, *p*=0.0862, Fig A2a) and Tp (*F*1,105 =0.3899, *p*=0.760, Fig A2b).

### 3.3 Ecophysiological traits

Increasing temperatures caused decreases in fat (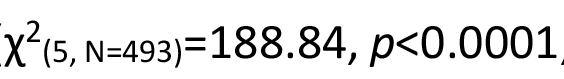, Fig 5a), protein (*F*5,478 =9.88, *p*<0.0001, Fig 5b) and sugar content (*F*5,460 =18.942, *p*<0.0001, Fig 5c). Overall, Senegalese snails had a higher protein content than snails from the other countries (*F*3,478 =7.10; *p*<0.001, Fig 5a, all contrasts: *p*<0.0326). Fat (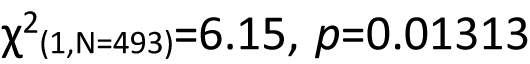, Fig 5a) and sugar (*F*3,461=4.562, *p*=0.033, Fig 5c) contents decreased as the number of weeks survived increased. Furthermore, the fat (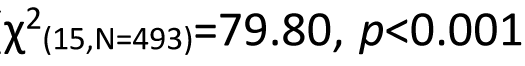, Fig 5a) and sugar (*F*15,460=3.609, p<0.001, Fig 5c content were affected by temperature and this differed across countries. The Malilangwe population had both the lowest fat content, but only at 12 °C (contrasts: all *p*≤0.0035) and 32 °C (contrasts: all *p*≤0.0190), and the lowest sugar content, but only at 8 °C (contrasts: all *p*≤0.0233) and 12 °C (contrasts: all *p*≤0.001). French snails on the other hand had the highest sugar contents than snails from all other countries at 8 °C (contrasts: all *p*≤0.0365; *p*=0.001, Fig 5c) and compared to snails from Malilangwe at 12 °C (contrast: *p*<0.001). At 32 °C, French snails had a lower fat content than Senegalese (contrast: *p*=0.0004), and Zimbabwean (contrast: *p*=0.0190) snails, yet a higher fat content than snails from Malilangwe (contrast: *p*=0.0190). At the other temperatures, there were no differences in fat content (contrasts: all *p*>0.0935) nor in sugar content (contrasts: all *p*>0.1337) compared to the other countries. No interaction effect between temperature and country was found for the protein content (*F*15,464, *p*=0.5688, Fig 5b).

**Figure 5:**
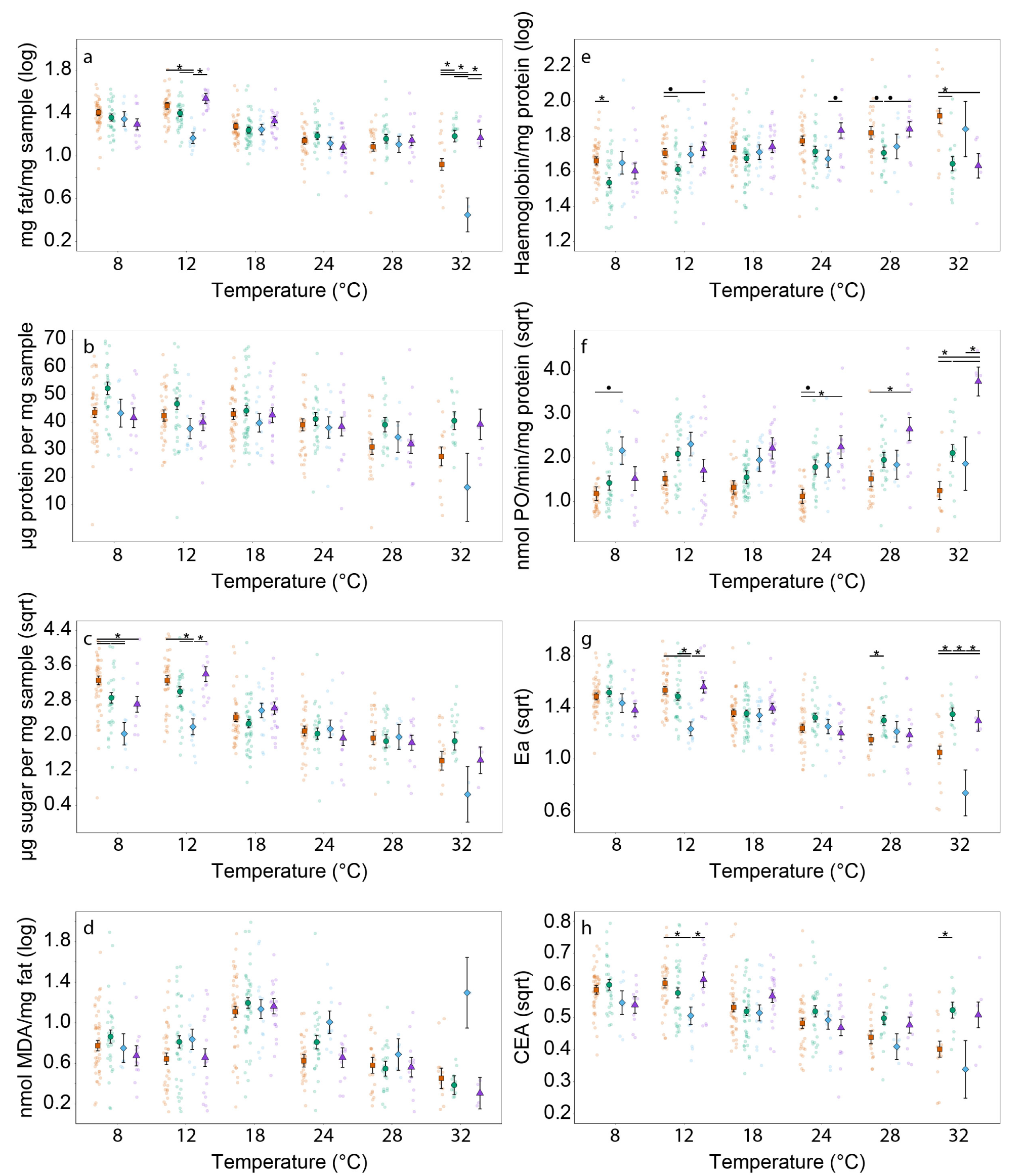
Results of the ecophysiological analyses. a) fat content, b) protein content, c) sugar content, d), oxidised fat (MDA), e) haemoglobin, f) phenoloxidase (PO) levels, g) total energy available (Ea), and h) total energy budget (CEA) as a function of temperature and country of origin. Significant (*p*<0.05) interaction effects of temperature and country of origin are indicated by asterisks. Trends are indicated by bullets.

**Figure 6.**
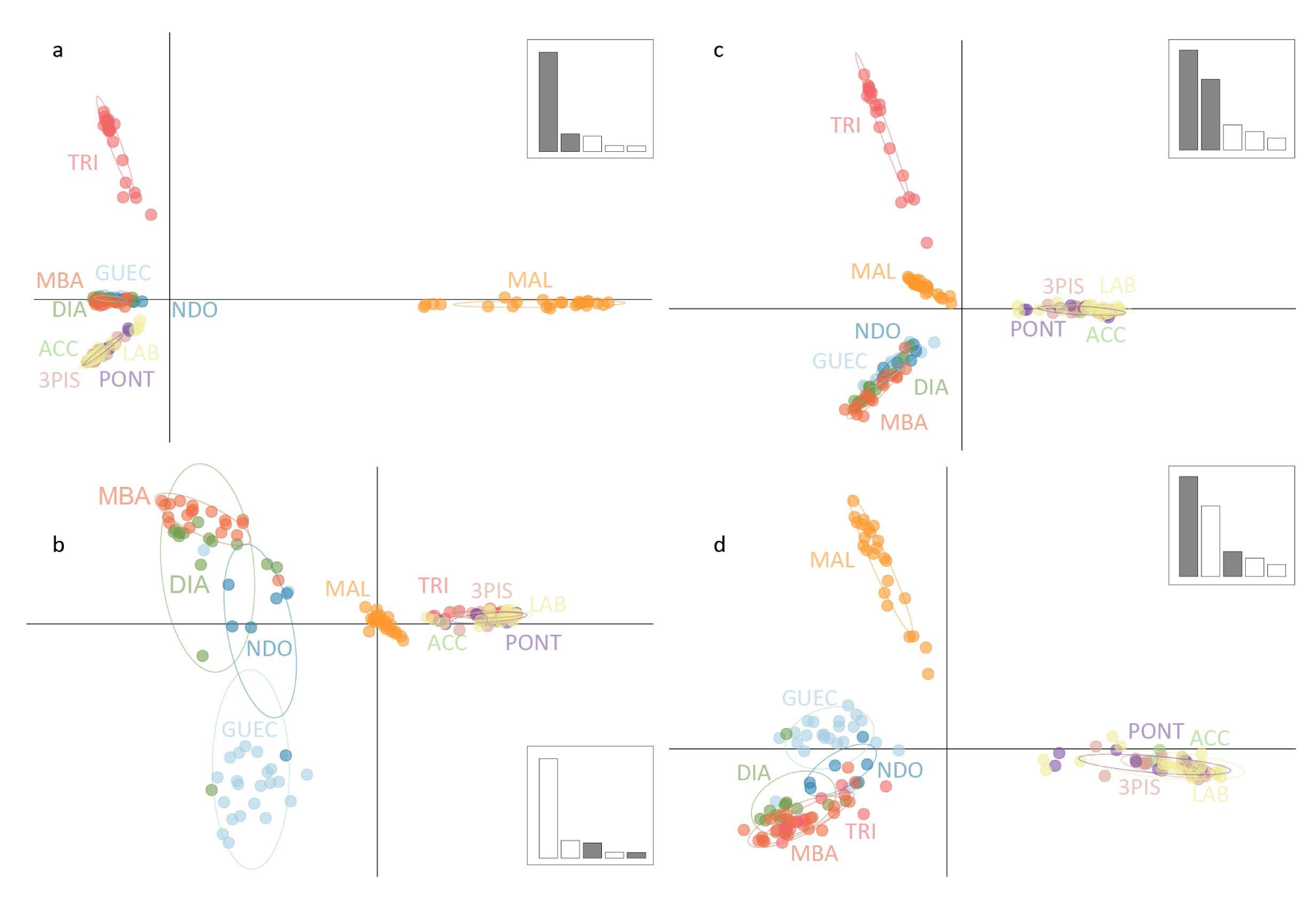
The PCA analyses of the neutral (a, b) and outlier (c,d) datasets. a): PCA axes 1 and 2 and b) PCA axes 3 and 5 of the neutral dataset; c) PCA axes 1 and 2 and d) PCA axes 1 and 3 for the outlier dataset. Different colours indicate different sampling sites. The full names of the abbreviations can be found in Table 1.

Metabolic rate (ETS activity) increased with increasing temperatures (*F*5,475=24.76; *p*<0.001, Fig 5d). However, no significant differences in metabolic rate were found between countries (*F*3,475=1.37; *p*=0.250, Fig A3a).

Malondialdehyde (MDA) levels differed between countries (*F*3,476=4.707; *p*=0.003, Fig 5d), with MDA levels being higher in Malilangwe snails compared to French (contrast: *p*=0.0075) and Zimbabwean snails (contrast: *p*=0.0056) and tended to be higher than Senegalese snails (contrast: *p*=0.0957). Additionally, temperature affected MDA levels (*F*5,476=37.949; *p*<0.001, Fig 5e): MDA levels were highest at 18 °C and declined towards both higher and lower temperatures (all contrasts: *p*<0.001, Fig 5d). No interaction between country and temperature was observed (*F*15,461=1.1233, *p*=0.332).

Regarding the haemoglobin levels, the effect of temperature tended to depend on the country of origin (*F*15,460=1.56; *p*=0.0813, Fig 5e). The haemoglobin levels increased with increasing temperatures (contrasts: all *p*≤0.0438) but declined again at temperatures above 28 °C in Senegalese and Zimbabwean snails, however, haemoglobin levels kept on increasing in French and Malilangwe snails (*F*15,460=1.56; *p*=0.0806, Fig 5e). Overall, there was a difference between countries (*F*3,460=15.13, *p*<0.001, Fig 5e) with French (contrast: *p*<0.001) and Zimbabwean (contrast: *p*=0.0024) snails having higher haemoglobin contents than Senegalese snails (other contrasts: all *p*>0.1955).

Overall, French snails had the lowest and Zimbabwean snails had the highest PO levels (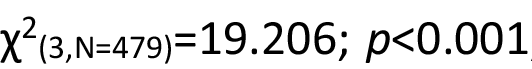, Fig 5f), although the post hoc tests lacked the power to detect these differences. While PO activity increased with temperature 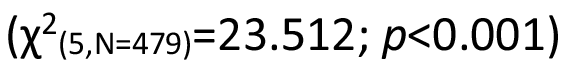, this effect differed across the countries of origin 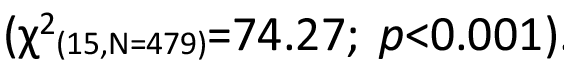. Specifically, Zimbabwean snails had the highest PO levels at 32 °C compared to French (contrast: *p*<0.001), Senegalese (contrast: *p*=0.0021) and Malilangwe (contrast: *p*=0.0403) snails, while French snails had lower PO levels than Senegalese snails at this temperature (contrast: *p*=0.0283). At other temperatures, no significant differences between the countries were observed (contrasts: all *p*≥0.0611).

The country of origin (*F*3,454=10.011, *p*<0.001) and the temperature (*F*5,454=21.827, *p*<0.001) significantly affected the energy available (Ea, Fig 5g). Furthermore, there was a significant interaction between the two (*F*15,454=3.627, *p*>0.001). The Malilangwe snails had less energy available than French, Senegalese, and Zimbabwean snails at 12 °C (all contrasts: *p*<0.001). Senegalese snails had more energy available than French snails at 28 °C (contrast: *p*=0.0345) and than French (contrast: *p*<0.001), Zimbabwean (contrast: *p*=0.0457), and Malilangwe (contrast: *p*=0.0057) snails at 32 °C. Finally, Zimbabwean snails had more energy available than Malilangwe snails at 32 °C (contrast: *p*=0.0226). No effects of temperature (*F*5,465=1.448, *p*=0.2057) or country of origin (*F*3,465=0.1911, *p*=0.9025) on the energy consumption (Ec) were detected (Fig A3b). The total cellular energy budget available (CEA) was affected by temperature (*F*5,454=15.19; *p*<0.001, Fig 5h), the country of origin (*F*3,454=5.016; *p*=0.00198, Fig 5h), and their interaction (*F*15,454=2.354; *p*=0.00291, Fig 5h). The highest energy budgets were observed in the cold temperature treatments (8-18 °C, contrasts: *p*≤0.0387) with a continuous decline towards the higher temperatures. Overall, Senegalese snails had a higher energy budget than French (contrast: *p*=0.0220) and Malilangwe (contrast: *p*=0.0092) snails.

### 3.4 Genetic diversity, structure and outliers

After SNP filtering, a total number of 12,875 polymorphic sites were discovered in 166 individuals. Bayescan identified a total of 1765 outlier SNP loci potentially subjected to adaptive selection. PCadapt was less selective and identified a total of 4,625 outlier SNPs, of which 633 were shared with Bayescan. These 633 loci were included in the outlier dataset while the remaining 12,242 loci constituted the neutral dataset.

Allelic richness values across populations ranged from 1.006 (Malilangwe) to 1.501 (Accrobranche) for the neutral SNP dataset and from 1.000 (Malilangwe) to 1.225 (Nombo) for the outlier dataset (Table A2). In both datasets, the Malilangwe population had a very low allelic richness, with only one allele at most loci. Pairwise *FST* values ranged from 0.0113 between Pont Mulinu and Laboratory to 0.664 between Malilangwe and Ndombo in the neutral dataset (Table A4), and from 0.0162 (3 Piscines and Accrobranche) to 0.905 (Malilangwe and Triangle) for the outlier dataset (Fig A5).

Regarding the neutral dataset, the PCA of axes 1 and 2 (explaining 40.53 and 7.22 % of the variance, resp.) allowed the resolution of four main groups corresponding to the three countries and the Malilangwe population clustering apart from the other populations along axis 1 (Fig 6a). PCA axes 3 and 5 (explaining 6.24 and 2.28 % of the total variance, resp.) allowed resolving the population structure of the Senegalese populations with the Guédé Chantier population clustering apart from the other Senegalese populations (Fig 6b). Regarding the outlier dataset, PCA axes 1 and 2 (explaining 31.30 and 22.16 % of the total variance, resp.) identified the same four main clusters as the neutral dataset but with a central clustering of the Malilangwe population (Fig 6c). PCA axes 1 and 3 (explaining 31.30 and 7.83% of the total variance, resp.) were able to identify the Guédé Chantier population as genetically different from the other Senegalese populations (Fig 6d). The STRUCTURE analysis on the neutral dataset showed Δ *K* values peaking at *K*=5 and supports the conclusion from the PCA analysis that the main hierarchical level of the population structure is based on five groups (Fig A6). Little allele sharing between the groups has been observed, except for the Senegalese Guédé Chantier and Ndombo populations at *K*=5 and *K*=6. Increasing *K* to seven did not provide further sample resolution (Fig A6).

Out of the 633 outlier loci, 165 could be associated with at least one environmental variable in the LFMM analysis (Table A3). Of these, 68 could be associated with precipitation, 76 with temperature, and 21 with both temperature and precipitation. The multivariate population-based RDA analysis on the complete and outlier dataset could not detect any significant association between the SNPs and environmental variables (for the complete dataset: *F*=0.619, 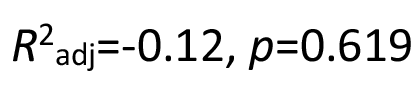; and for the outlier dataset: *F*=0.6272, 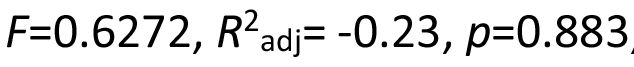, Fig A7). These findings were similar when other environmental variables and longitude/latitude coordinates were included in the analyses.

The RDA analysis on the outlier dataset to associate SNPs with the life history data was marginally non-significant 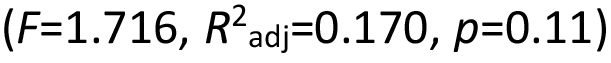. The backward selection procedure identified growth rates at 8 and 28 °C to be most plausibly related to some SNPs (Fig A8).

## 4 Discussion

Local adaptation to temperature is commonly observed across a wide range of species, including aquatic invertebrates (e.g. in *Daphnia* sp. (Yampolsky et al., 2013), crustaceans (Seefeldt & Ebert, 2019), and gastropods (Gleason & Burton, 2013)). However, the question remains if these adaptations are genetically based and could be driven causally by climate change (Stoks et al., 2014). We discovered trade-offs between life history and different ecophysiological traits in response to temperature in snail *B. truncatus* populations from three different climatic zones. Furthermore, we found some indications of genetic adaptation to climatic factors such as temperature and rainfall, providing the first evidence of a causal link between genetic variation and phenotypic traits that could drive climate adaptation.

### 4.1 A life history and ecophysiology adapted to local climate

*Bulinus truncatus* showed strong phenological and ecophysiological adaptations to temperature. We observed a marked trade-off between investment in survival, growth, and reproduction between the different climatic origins. In this respect, cold-origin snails have higher growth rates and allocate less energy to reproduction while warm-origin snails have higher reproduction rates that compensate for lower survival rates.

Growth rates increased with temperature for all countries, a pattern also observed in freshwater apple snails, *Pomacea canaliculate* and *Asolene platae* (Seuffert & Martín, 2013; Tiecher et al., 2015). Compared to other countries, the cold-origin (French) snails had relatively higher growth rates at both low and high temperatures that allow for efficient use of the short growing season to build up sufficient energy reserves to survive the cold winter (Dillon, 2000). Indeed, seasonal length is an important mechanism driving adaptive patterns (e.g. in Atlantic sturgeon and silverside fish: Baumann & Conover, 2011; Markin & Secor, 2020, and Swedish common frogs: Ståhlberg et al., 2001) and has also been observed in the marine snail *Urosalpinx cinerea*, which has high growth rates early in the season while delaying spawning to benefit from warmer temperatures later in the season (Villeneuve et al., 2021). The larger final size of French snails follows macroecological patterns such as James’ rule and Bergmann’s rule, which state that within species, larger individuals are found at higher, colder latitudes (James, 1970; Van Voorhies, 1996). The decreasing size of ectothermic organisms at lower latitudes could be attributed to the reduced availability of oxygen (Rollinson & Rowe, 2018). Although *B. truncatus* experiences temperatures as low as 4 °C for a couple of weeks in the Cavu river in Corsica, France (Mulero et al., 2019), all snails in our study were dead after one week at this temperature. This is in stark contrast to the study by Mulero et al. (2019) who found that cold-origin *B. truncatus* survive over 15 weeks at 4 °C. However, in their study, the temperature was decreased from 25 °C to 4 °C over 40 days, compared to 10 days in this study. This suggests that the snails require more time to acclimate and lower their cold tolerance through adaptive plasticity (as observed by Loomis (1985) in the freshwater snail *Melampus bidentatus*).

While reproduction is positively correlated with temperature in all countries, warm-origin (Senegalese) snails allocate a greater proportion of their available energy to reproduction at higher temperatures compared to the other countries (both in terms of the number of egg masses laid and the number of eggs per egg mass). This causes a trade-off that results in reduced growth and survival under elevated temperatures. Indeed, the Senegalese hot season, spanning from July until October, with a mean water temperature of 29.7 °C in the Senegal River (Ernould et al., 1999), coincides with a sharp decline in the abundance of *B. truncatus* snails (Ndione et al., 2018). The high reproduction rate exhibited at these temperatures compensates for the lower survival rates and acts as a protective mechanism against population extinctions (Conover et al., 2009; Daufresne et al., 2009). Furthermore, producing a high number of offspring can be a bet-hedging strategy that increases the population’s fitness under heat stress (Sergio et al., 2018) or a response to other selective pressures such as predation (Lips, 2001). This trade-off between survival/growth and reproduction is also present in various other species, such as spider mites (Li et al., 2022), fruit flies (Marshall & Sinclair, 2010), and land snails (Çelik et al., 2022). Finally, adaptation to acute thermal stress (higher CTmax values) allows the warm-origin snails to move to cooler (shaded or deeper) areas when the water is directly exposed to the Sahelian sun and the top water layer warms up quickly (Kuo & Sanford, 2009).

Our findings only partially align with the results of Konan et al. (2022) and Diakité et al. (2023) who revealed variation in the life-history traits of *B. truncatus* between one warm-origin population from Northern Ivory Coast and one cold-origin population from Central Ivory Coast (but both stemming from a tropical savanna climate with a dry winter). Similarly to our study, the warm-origin population attained an earlier age of reproduction and had higher egg-laying rates at 24 °C than the cold-origin population. However, contrary to our results, they report higher survival rates in the warm-origin population at 24 °C with up to 60 % of the snails still surviving after 20 weeks (in comparison to the 10 % survival at 24 °C for the warm-origin snails in our study). A noteworthy disparity between our study and theirs lies in the fact that they studied only two snail populations originating in one ecoregion with smaller climatic differences, while our research is based on genotyped snails from multiple populations and ecoregions, ruling out cryptic diversity and limiting the chances of sampling artefacts. Additionally, they used the first generation offspring which might be more subject to maternal effects.

More subtle patterns of local adaptation are observed in the ecophysiological responses. At high temperatures, more energy is allocated towards growth and reproduction which causes the net energy budget (CEA levels) to decrease with increasing temperatures. This is mainly attributable to the declining energy reserves (Ea levels, driven by decreasing protein, sugar and fat reserves), although the rates differ between countries (as observed in damselflies in Verheyen et al., 2023). Cold-origin snails had a higher fat and sugar content at 12 °C, which results in a higher CEA and is probably associated with the lower energy investment in reproduction (lower number of egg masses and eggs per egg mass at lower temperatures). This energy can be invested in locomotor performance and metabolic maintenance as has been observed in the apple snail *Pomacea canaliculata* (Matsukura et al., 2009).

The elevated protein contents of the warm-origin snails potentially indicate heightened levels of heat shock proteins to safeguard against protein denaturation, or elevated antioxidant defence mechanisms to avoid oxidative stress and damage (Clusella-Trullas et al., 2013; Jeyachandran et al., 2023; Wang et al., 2022). Nevertheless, Senegalese snails still experienced increased levels of oxidative damage to lipids (i.e. MDA levels), implying that the antioxidant defence system may not efficiently eliminate the generated reactive oxygen species (ROS) promptly (Monaghan et al., 2009; Oksala et al., 2014). This may result in the reduced survival rates of populations from this country at higher temperatures. Additionally, the Senegalese snails had a lower haemoglobin content, potentially because of their enhanced oxygen-binding capacity of haemoglobin, resulting from living in warm, oxygen-deficient waters (Bugge & Weber, 1999).

Notably, the Zimbabwean snails had the highest phenoloxidase (PO) levels and appeared to prioritise investments in immune defence over rapid growth or high egg-laying rates (as observed in damselflies: De Block & Stoks, 2008). The high PO levels can be attributable to the higher parasite diversity in Zimbabwe as compared to France and Senegal (Pappalardo et al., 2020; Salkeld et al., 2008; Thieltges et al., 2011). Indeed, having high PO levels when living in parasite-rich areas (i.e. Zimbabwe) is important as PO strongly contributes to the immune defence against trematodes and other parasitic infections in snails (Le Clec’h et al., 2016). Additionally, just as in flies (Gourgoulianni et al., 2023), PO levels in *B. truncatus* increase with temperature. This increase in PO can be attributed to a higher pathogen growth (Harvell et al., 2002) or the increased metabolic activity of ectotherms (Clark & Worland, 2008) at higher temperatures. However, it should be noted that PO levels are not always positively correlated with temperature as shown in the great pond snail *Lymnaea stagnalis* (Seppälä & Jokela, 2011).

**4.2 Genetic basis for local adaptation in *Bulinus truncatus***

In addition to the patterns observed in the life history and ecophysiology, we were able to detect genetic signs of local adaptation in *B. truncatus* unconfounded by any maternal effects or phenotypic plasticity. We identified five well-delineated genetic clusters with the populations from each country clustering apart and two separate clusters in both Senegal (the Guédé-Chantier population clustering apart from Ndombo, Mbane and Diama) and Zimbabwe (Malilangwe and Triangle) and limited gene flow between populations. This high level of population structuring is comparable to previous studies on *B. truncatus* (Chlyeh et al., 2002; Maes et al., 2022; Zein-Eddine et al., 2017) and results in a high potential for genetic adaptation as limited gene flow between populations is a prerequisite to maintaining specialised, locally adapted, genotypes (Blanquart et al., 2013, but see Tigano & Friesen, 2016).

The percentage of outliers detected in our study bears resemblance to findings in another study on a marine snail species (Simmonds et al., 2020), with a substantial proportion (26.07%) of these outliers demonstrating potential correlations with temperature, precipitation, or both, illustrating the highly polygenic nature of adaptation to climate. However, the position of these SNPs on the genome could not be determined since no fully assembled reference genome for *B. truncatus* is available (Young et al., 2022). It is worth noting that the actual number of genome-wide SNPs associated with environmental factors is expected to be considerably higher, given that our study only sampled a small fraction of the genome using GBS, suggesting a significant potential for local adaptation within the *B. truncatus* genome. The limited available landscape genomic studies on freshwater species support our findings as they indicate that genetic adaptation to the environment is indeed feasible, even in highly interconnected and mobile freshwater species (such as in fish cf. Deflem et al., 2022 and Harrisson et al., 2017).

The non-significance of the RDA to test for an association between SNPs and climatic variables is likely attributable to limited statistical power and data structure. The inclusion of only eight populations from three different countries does not adequately represent a full gradient in environmental conditions. Localities within each country exhibit similar climatic conditions, effectively resulting in only three distinct climate categories to which the SNPs could be linked. This is also reflected in the RDA analysis of the life history data (genotype-phenotype analysis). The inclusion of eight populations and the population’s specific life history data substantially enhances the statistical power of the test, revealing some SNPs that were correlated with the growth rate of the snails, although also this test was non-significant.

The notable genetic segregation of the Malilangwe population from all other groups, including the nearby Triangle population that is situated approximately 80 kilometres away, could probably be attributed to two key factors. Firstly, the relatively low allelic richness observed in the Malilangwe population may be attributed to a pronounced founder effect, as documented by Nalugwa et al. (2011). This population was collected from an artificial lake established in 1964 within a nature reserve. When the lake was filled, the local riverine snail population may have diminished, giving rise to a new population potentially formed through intensive self-fertilisation of a limited number of surviving aphallic individuals (Brown, 1994; Jarne et al., 1994). In contrast, the Triangle site represents a temporary lake within the same river system characterised by substantial anthropogenic influence and frequent cattle visits, potentially promoting gene flow from nearby snail populations (Kappes & Haase, 2012). Moreover, the recurrent cycles of partial drying and refilling of this lake could contribute to the increased genetic diversity, as various populations mix in the deepest parts of the lake during drying phases (Viard et al., 1996, 1997). Secondly, it is crucial to acknowledge that *B. truncatus* is part of the *Bulinus truncatus/tropicus* complex, a group of closely related species that pose significant challenges in both genetic and morphological identification (Brown, 1994; Brown & Shaw, 1989; Nalugwa et al., 2010), with individuals from Malilangwe already possessing some level of genetic divergence. A more comprehensive investigation, including a more detailed assessment of ploidy levels and an examination of the radula and reproductive organs (Brown, 1994), may provide further insights.

### 4.3 Implications for schistosomiasis transmission

Local adaptation in vector populations plays a profound role in shaping the dynamics of schistosomiasis transmission in various ways. Firstly, the effect of climate change on the distribution of schistosomiasis mainly depends on the tolerance limits, dispersal and adaptation capacity of the host snails. Schistosome parasites exhibit broader tolerance ranges than their snail hosts (Mulero et al., 2019), and they can easily be introduced in new areas by infected people (e.g. the introduction of a Senegalese schistosome strain in Corsica by infected tourists; Boissier et al., 2016; Moné et al., 2015). Therefore, schistosomes introduced in non-native areas where a compatible *B. truncatus* population is present, can readily establish irrespective of the environmental conditions. Additionally, the high dispersal capacity of snails, facilitated by animal and human vectors (Kappes & Haase, 2012; Pfenninger et al., 2011), enables them to traverse barriers such as seas or mountain ranges. Consequently, the northward spread of schistosomiasis in Europe is currently constrained by the tolerance limits of *B. truncatus* to colder temperatures. Our study showed that these tolerance limits are subject to local adaptation in *B. truncatus*, potentially altering them within a few generations, as demonstrated in the case of the mosquito *Aedes aegypti* (Dennington et al., 2024). This adaptability could potentially expand the distribution area of the species over time. Therefore, incorporating the adaptation capacity of the intermediate host snails into assessments of the impact of climate change on schistosomiasis risk could increase the reliability of these predictions (DeMarche et al., 2019; Valladares et al., 2014).

Secondly, vector life-history traits are a crucial determinant of human infection dynamics in multiple vector-borne diseases. For example, in mosquito-borne diseases, variations in vector characteristics like biting rates, reproduction rates, and development times result in distinct transmission rates to humans (Mordecai et al., 2019). Similarly, the higher available energy budget (in terms of sugar and fat content) of the cold-origin snails might result in more and bigger parasites because more energy is available for schistosome development (Mas-Coma et al., 2009). Additionally, the high reproductive capacities of the warm-origin snail populations found in our study generate highly age-stratified populations that can result in an elevated parasite output (Anderson et al., 2021). The pronounced age-structuring of Senegalese *B. truncatus* snails and the associated high parasite output provides a partial explanation for the sustained prevalence of urogenital schistosomiasis in the Senegal River Basin, despite sustained control efforts (Kokaliaris et al., 2021).

Finally, the adaptive capacity of *B. truncatus* demonstrated in this study may confer increased resilience to snail control efforts. Like other disease vectors (e.g. mosquitoes; Smith et al., 2016), *B. truncatus* could evolve resistance to chemical control agents (Konan et al., 2022), potentially diminishing their effectiveness. Furthermore, local adaptation could also manifest in response to predator cues (Dalesman et al., 2015; Goeppner et al., 2020). This implies that biological control measures reliant on the introduction of snail predators, such as fish (Arostegui et al., 2019), waterbugs (Younes et al., 2017), or prawns (Faiad et al., 2023; Monde et al., 2017; Swartz et al., 2015), may exhibit reduced efficacy in areas where natural predators are already present.

## 5 Conclusions

*Bulinus truncatus* shows a significant potential to adapt to its local environment on a morphological, ecophysiological and genetic level. However, further research is imperative to comprehensively evaluate the local adaptation potential to various other biotic and abiotic environmental factors. Specifically, future investigations should delve deeper into elucidating the mechanisms underlying local adaptation and the consequences it holds for the response of *B. truncatus* to a changing climate. Furthermore, the adaptation potential of *B. truncatus* implies that this species could adapt to chemical or biological snail control measures. These insights bear significance for prospective endeavours aimed at predicting the future distribution of intermediate hosts and, by extension, the distribution of schistosomiasis under global change, but also for the design of effective snail control efforts in the context of schistosomiasis elimination.

### CRediT authorship contribution statement

**Tim Maes:** Conceptualisation, Methodology, Formal analysis, Funding acquisition, Investigation, Project administration, Visualisation, Writing-original draft, Writing-review & editing. **Julie Verheyen:** Methodology, Formal analysis, Validation, Writing-original draft, Writing-review & editing. **Bruno Senghor:** Resources, Writing-review & editing. **Aspire Mudavanhu:** Resources, Writing-review & editing, **Bart Hellemans:** Formal analysis, Investigation, Resources. **Enora Geslain:** Formal analysis, Software, Methodology. **Ruben Schols:** Resources, Writing-review & editing. **Filip A.M. Volckaert:** Conceptualisation, Methodology, Supervision, Funding acquisition, Writing-review & editing. **Hugo F. Gante:** Methodology, Supervision, Writing-review & editing. **Tine Huyse:** Conceptualisation, Methodology, Supervision, Funding acquisition, Writing-review & editing.

## Acknowledgements

Tim Maes benefited from an FWO fellowship grant (ref. no. 1S86319N) of the Research Foundation—Flanders and travel grants from the FWO (ref. no. K203921N) and Royal Belgian Zoological Society (RBZS) for the snail collections in Senegal. Julie Verheyen is a postdoctoral fellow of the FWO (ref. no. 12ZZX21N). Ruben Schols was supported by BRAIN-be 2.0 under the MicroResist project (Grant number B2/191/P1/MicroResist). Aspire Mudavanhu was supported financially by the Global Minds Ph.D Fellowship of the Flemish Interuniversity Council - University Development Co-operation (VLIR-UOS) and logistically by the Malilangwe Trust. Both Ruben Schols and Aspire Mudavanhu benefited from a travel grand from the Stichting ter Bevordering van het Biodiversiteitsonderzoek in Afrika (SBBOA) for the snail collections in Zimbabwe. The authors thank the field teams in Senegal and Zimbabwe for assisting in the snail collections and Jérôme Boissier from the University of Perpignan Via Domitia for supplying the Corsican snail strains. We thank Ria Van Houdt for analytical support, Geert Neyens and Rony Van Aerschot for technical support during the experiment, and Yves Bawin for bio-informatic advice. Some of the research was carried out with infrastructure funded by the European Marine Biological Resource Centre (EMBRC) Belgium, Research Foundation - Flanders (FWO) project (ref. no. I001621N).

