## Supplementary material for "First evidence of a causal link between genetic variation and thermal adaptation in a schistosome host snail"

**Appendix 1 detailed protocols physiology**

Protein concentration of the homogenate was measured using the Bradford (1976) method while fat content was measured using a modified protocol of Marsh and Weinstein (1966). For the fat content, glass tubes were filled with 8 µl homogenate and 56 µl sulphuric acid (100 %), and heated at 150 °C for 20 min. The tubes were cooled down and 64 µl Milli-Q water was added. From this mixture, 30 µl was added to a 384-well plate in triplicates per sample and absorbance was measured at 490 nm. A standard curve of glyceryl tripalmitate was used to convert the absorbance into fat content (using the mean of the three readings). Fat content was expressed in µg per mg wet weight. The total sugar content (glucose + glycogen) was determined by adding 5 µl homogenate to 13 µl milliQ water and 2 µl amyloglucosidase (Sigma A7420; conc. 1 unit/10 µl milliQ water) in a 384-well plate. The mixture was incubated for 30 minutes at 37 °C after which 40 µl glucose reagent (Sigma G3293) was added. The samples were again incubated for 20 minutes at 30 °C and the absorbance was measured at 340 nm in duplicate. A standard curve of glucose was used to convert the absorbance into total sugar content (using the mean of the two readings).

To measure the haemoglobin content (Lieb et al., 2006), 40 µl sample homogenate was added to a 384-well plate and the absorbance was measured at wavelengths of 560, 576 and 600 nm. Haemoglobin contents were calculated as $[A_{576}-(A_{560}-A_{600})/2]$ x dilution factor/(mg protein/wet mass) (cf. Yampolsky et al., 2013).

The level of malondialdehyde (MDA) was measured using a modified protocol of Miyamoto et al. (2012) to estimate the oxidative damage to lipids. First, 25 µl homogenate was further diluted with 25 µl PBS buffer and 50 µl 2-Thiobarbituric acid (TBA) solution (0.4 % in 0.1 M HCl). This solution was then incubated at 100 °C for 60 minutes. After cooling, 165 µl butanol was added and mixed vigorously. The mixture was then centrifuged for 3 minutes at 4000 g. Finally, 30 µl of the mixture was transferred to a 384-well plate in triplo and fluorescence was measured at an excitation/emission wavelength of 535/550 nm. The standard curve of 1,1,3,3-tetramethoxypropan 99 % malonaldehyde bis (dimethyl acetol) 99 % was used to calculate MDA concentrations expressed as nmol MDA per mg fat.

Phenoloxidase (PO) activity was used as a measure of immune function and was quantified using a modified protocol of Stoks et al. (2006). In freshwater snails, PO has been associated with the immune response against trematode infections (Le Clec'h et al., 2016). We added 15 µl of the sample homogenate, together with 5 µl chymotrypsin (1 mg/ml Milli-Q water) in a 384-well microtitre plate (in duplo). After incubating for 5 minutes at room temperature to allow the conversion of proPO to PO, 20 µl L-Dopa (1.97 mg/ml PBS) was added. Phenoloxidase catalyses the transition from L-Dopa to dopachrome and this reaction proceeded at 30 °C for 30 minutes while absorbance of dopachrome was measured every 20 s at 490 nm. The slope of the linear part of the reaction curve (time interval 1500 – 6000 s) was used to quantify the PO activity. The average of the duplicate readings per sample was used for statistical analyses and one unit of PO activity is expressed in nmol dopachrome formed per minute per mg protein.

The metabolic rate of the snails was quantified at the cellular level by assessing the ETS activity as measured by De Coen and Janssen (2003). In a 384-well plate, 5 µl of the homogenate was mixed with 15 µl buffered substrate solution (0.13 M Tris-HCl, 0.3 % triton X-100, 1.7 mM NADH, 250 µM NADPH, pH 8.5). Next, 10 µl iodonitrotetrazolium (INT, 8 mM p-iodonitrotetrazolium) was added. The INT replaces $O_2$ as an electron acceptor and receives electrons from NADPH via NADH-cytochrome oxidoreductase with the formation of formazan as a result. The increase in absorbance was measured in duplo, every 30 s for 30 minutes at 490 nm and 20 °C. The formazan concentrations were calculated based on the Lambert-Beer law using a molecular extinction coefficient of 15.9 $mM^{-1}cm^{-1}$. The observed concentrations were then converted to cellular oxygen consumption rates using the stoichiometric relationship that 1 µmol $O_2$ is used to form 2 µmol formazan in the ETS system. ETS activity was expressed as nmol $O_2$ per minute per mg protein.

The cellular energy allocation (CEA) was calculated based on the total protein, fat and sugar content as outlined in Gomes et al. (2015). First, to calculate the total energy available (Ea), the protein, sugar and fat contents were converted to energy contents by multiplying the nutrient contents by the respective energy equivalents (24 kJ/g proteins, 17.5 kJ/g sugars, 39.5 kJ/g lipids). Next, the total energy consumption (Ec) was determined based on the values for the ETS measured. The amount of consumed oxygen is transformed into energetic equivalents using the specific oxyenthalpic equivalents for an average lipid, protein and carbohydrate mixture of 484 kJ/mol $O_2$. Finally the CEA is calculated as the ratio between Ea and Ec.

**Appendix 2 raw nQuire results**

| file | free | dip | tri | tet | d_dip | d_tri | d_tet |
|---|---|---|---|---|---|---|---|
| 10.bin | 38800.183557 | 9362.136430 | 23556.141919 | 34437.610131 | 29438.047127 | 15244.041638 | 4362.573426 |
| 100.bin | 33781.047506 | 3295.014637 | 15643.513285 | 26795.672782 | 30486.032870 | 18137.534221 | 6985.374724 |
| 101.bin | 52013.181161 | 8926.831867 | 27468.295663 | 42954.159995 | 43086.349294 | 24544.885498 | 9059.021166 |
| 102.bin | 61665.420096 | 12947.484959 | 35680.191676 | 53224.617749 | 48717.935138 | 25985.228421 | 8440.802347 |
| 103.bin | 55450.509260 | 14731.959203 | 33429.442729 | 46852.740784 | 40718.550058 | 22021.066532 | 8597.768476 |
| 104.bin | 8630.339129 | -267.416694 | 2310.754796 | 5368.564641 | 8897.755823 | 6319.584333 | 3261.774488 |
| 105.bin | 925.575345 | -102.463174 | 176.278877 | 568.921556 | 1028.038519 | 749.296468 | 356.653789 |
| 107.bin | 53100.643931 | 13175.399277 | 29651.309857 | 43096.188545 | 39925.244654 | 23449.334074 | 10004.455386 |
| 108.bin | 6112.414807 | 2267.478917 | 4044.062846 | 5222.365252 | 3844.935890 | 2068.351962 | 890.049555 |
| 109.bin | 58265.352471 | 14605.278412 | 33723.553940 | 48209.359349 | 43660.074059 | 24541.798531 | 10055.993122 |
| 11.bin | 47622.624541 | 12932.253777 | 30113.279790 | 42313.774373 | 34690.370764 | 17509.344750 | 5308.850167 |
| 110.bin | 75522.370810 | 39363.024533 | 52282.617664 | 62783.540004 | 36159.346276 | 23239.753146 | 12738.830805 |
| 111.bin | 36766.548943 | 10781.955956 | 23964.598264 | 33218.590479 | 25984.592987 | 12801.950679 | 3547.958464 |
| 112.bin | 65732.772534 | 25996.093836 | 40652.508037 | 52218.360845 | 39736.678698 | 25080.264497 | 13514.411689 |
| 113.bin | 72099.476369 | 37008.306098 | 49806.709249 | 59525.418022 | 35091.170271 | 22292.767120 | 12574.058346 |
| 114.bin | 60620.273140 | 20579.753847 | 36863.205275 | 48861.582265 | 40040.519292 | 23757.067865 | 11758.690875 |
| 115.bin | 62616.627937 | 23415.885542 | 40362.854670 | 52185.568441 | 39200.742394 | 22253.773267 | 10431.059495 |
| 116.bin | 67223.075814 | 21947.221609 | 43306.788746 | 57907.371204 | 45275.854205 | 23916.287068 | 9315.704610 |
| 117.bin | 45397.381469 | 6381.724579 | 22543.921363 | 37062.645462 | 39015.656890 | 22853.460107 | 8334.736007 |
| 118.bin | 66810.582155 | 30123.627975 | 46120.436232 | 57555.470005 | 36686.954181 | 20690.145924 | 9255.112150 |
| 119.bin | 62666.747423 | 17086.242240 | 36484.798174 | 51083.460756 | 45580.505183 | 26181.949249 | 11583.286666 |

| | | | | | | | |
|---|---|---|---|---|---|---|---|
| 120.bin | 54732.130127 | 9789.385689 | 28218.558569 | 43957.812486 | 44942.744439 | 26513.571559 | 10774.317641 |
| 121.bin | 87598.096061 | 45126.733977 | 64167.435158 | 75694.756146 | 42471.362083 | 23430.660902 | 11903.339915 |
| 122.bin | 50895.129993 | 7672.946268 | 25436.455168 | 40605.447176 | 43222.183725 | 25458.674826 | 10289.682817 |
| 123.bin | 66537.558007 | 27828.778711 | 44359.648547 | 55934.463996 | 38708.779296 | 22177.909460 | 10603.094011 |
| 124.bin | 44369.296616 | 4548.480866 | 21024.805100 | 35908.168633 | 39820.815751 | 23344.491516 | 8461.127983 |
| 125.bin | 61495.373084 | 29814.072397 | 41328.411857 | 49960.731894 | 31681.300686 | 20166.961227 | 11534.641190 |
| 126.bin | 58342.633372 | 12300.053037 | 32083.451277 | 48200.734490 | 46042.580335 | 26259.182095 | 10141.898882 |
| 127.bin | 31733.809692 | 1364.649877 | 13196.542031 | 25103.965684 | 30369.159815 | 18537.267661 | 6629.844008 |
| 128.bin | 29172.210212 | -577.924792 | 9877.873667 | 21737.785299 | 29750.135004 | 19294.336545 | 7434.424913 |
| 129.bin | 29629.368940 | -2069.981871 | 7724.460174 | 19876.690515 | 31699.350811 | 21904.908766 | 9752.678425 |
| 12.bin | 66254.079220 | 25553.990374 | 42044.003075 | 55291.000265 | 40700.088845 | 24210.076144 | 10963.078955 |
| 130.bin | 30340.201201 | 4393.763883 | 15427.093427 | 25169.020427 | 25946.437317 | 14913.107773 | 5171.180774 |
| 131.bin | 31607.005033 | 5204.359908 | 16971.702520 | 26934.169182 | 26402.645126 | 14635.302513 | 4672.835851 |
| 132.bin | 41298.996230 | 9366.115553 | 24753.379279 | 36745.222269 | 31932.880677 | 16545.616951 | 4553.773961 |
| 133.bin | 12808.451870 | 2788.371837 | 7458.831066 | 11388.590943 | 10020.080033 | 5349.620804 | 1419.860927 |
| 134.bin | 32932.254769 | 5040.890567 | 17323.878217 | 28070.600179 | 27891.364202 | 15608.376552 | 4861.654591 |
| 135.bin | 65482.181393 | 24854.733661 | 41495.010049 | 54397.432514 | 40627.447732 | 23987.171344 | 11084.748880 |
| 136.bin | 19817.463913 | 4403.143510 | 11950.712877 | 17927.698305 | 15414.320403 | 7866.751036 | 1889.765608 |
| 137.bin | 64362.178596 | 31267.671110 | 43499.795195 | 53622.575067 | 33094.507486 | 20862.383401 | 10739.603529 |
| 138.bin | 52044.511339 | 10312.050346 | 28317.379084 | 43094.909199 | 41732.460993 | 23727.132256 | 8949.602140 |
| 139.bin | 58306.453118 | 14374.898595 | 33071.655583 | 47549.776533 | 43931.554523 | 25234.797535 | 10756.676585 |
| 13.bin | 64798.769955 | 20508.917254 | 40002.363584 | 54488.255643 | 44289.852701 | 24796.406371 | 10310.514312 |

| | | | | | | | |
|---|---|---|---|---|---|---|---|
| 140.bin | 66194.442822 | 30873.668585 | 43448.847405 | 54245.247621 | 35320.774237 | 22745.595417 | 11949.195202 |
| 141.bin | 54751.909210 | 11488.721625 | 30386.537558 | 45073.961871 | 43263.187585 | 24365.371652 | 9677.947339 |
| 142.bin | 29675.120362 | 8055.533196 | 19415.979576 | 26972.871431 | 21619.587166 | 10259.140786 | 2702.248931 |
| 143.bin | 17476.645286 | 1546.570797 | 8233.380425 | 14611.286551 | 15930.074490 | 9243.264861 | 2865.358735 |
| 144.bin | 2356.017881 | 683.779465 | 1399.737250 | 1934.610147 | 1672.238416 | 956.280631 | 421.407734 |
| 145.bin | 38932.222494 | 9168.634740 | 22161.701655 | 32046.771008 | 29763.587754 | 16770.520839 | 6885.451486 |
| 146.bin | 8844.526723 | -192.244540 | 2444.161008 | 5483.164610 | 9036.771263 | 6400.365715 | 3361.362112 |
| 147.bin | 63808.695947 | 32911.570886 | 41237.691206 | 50962.360994 | 30897.125060 | 22571.004741 | 12846.334953 |
| 148.bin | 61845.359600 | 18816.766575 | 39634.895254 | 54063.200580 | 43028.593026 | 22210.464346 | 7782.159021 |
| 149.bin | 76028.541004 | 44357.414934 | 55404.351930 | 64240.243470 | 31671.126070 | 20624.189075 | 11788.297534 |
| 14.bin | 58909.703485 | 14856.757600 | 34528.631866 | 49480.378512 | 44052.945885 | 24381.071618 | 9429.324973 |
| 150.bin | 53690.376043 | 9151.984647 | 27866.669134 | 43425.934062 | 44538.391396 | 25823.706908 | 10264.441981 |
| 151.bin | 65669.020738 | 33228.334020 | 44289.898851 | 54284.691672 | 32440.686718 | 21379.121886 | 11384.329066 |
| 152.bin | 67615.285588 | 27839.906286 | 42835.197937 | 55527.502377 | 39775.379302 | 24780.087651 | 12087.783211 |
| 153.bin | 40663.435274 | 4884.281309 | 19905.660667 | 32934.645090 | 35779.153965 | 20757.774607 | 7728.790185 |
| 154.bin | 51562.982533 | 11944.702313 | 30097.560480 | 43683.405385 | 39618.280219 | 21465.422052 | 7879.577148 |
| 155.bin | 63797.767392 | 30570.848616 | 42895.694441 | 52408.321250 | 33226.918777 | 20902.072951 | 11389.446142 |
| 156.bin | 45413.468875 | 9341.225352 | 25600.709478 | 38625.765022 | 36072.243523 | 19812.759397 | 6787.703853 |
| 157.bin | 49268.039039 | 11320.784220 | 29208.753850 | 43145.166219 | 37947.254819 | 20059.285189 | 6122.872820 |
| 158.bin | 55086.496449 | 8415.356853 | 27596.378473 | 43631.152492 | 46671.139596 | 27490.117976 | 11455.343956 |
| 159.bin | 60468.755228 | 29742.882255 | 39406.816869 | 49444.816195 | 30725.872973 | 21061.938360 | 11023.939033 |
| 15.bin | 47089.174376 | 5621.037212 | 23892.759317 | 39848.264164 | 41468.137164 | 23196.415059 | 7240.910212 |
| 160.bin | 55409.217124 | 15750.141147 | 33879.197394 | 46578.931627 | 39659.075977 | 21530.019730 | 8830.285497 |

| | | | | | | | |
|---|---|---|---|---|---|---|---|
| 161.bin | 47400.773280 | 8523.125515 | 25344.092879 | 39239.129426 | 38877.647766 | 22056.680402 | 8161.643854 |
| 163.bin | 72425.272162 | 38177.153246 | 48126.586081 | 58569.617690 | 34248.118916 | 24298.686081 | 13855.654473 |
| 164.bin | 74787.507961 | 38124.090566 | 51698.856651 | 62673.902648 | 36663.417395 | 23088.651309 | 12113.605313 |
| 165.bin | 49264.212689 | 7668.178951 | 25202.907982 | 39873.662986 | 41596.033738 | 24061.304706 | 9390.549703 |
| 166.bin | 32884.504197 | 3484.626718 | 15508.839387 | 27037.913260 | 29399.877479 | 17375.664810 | 5846.590937 |
| 167.bin | 42080.401719 | 8254.035239 | 23877.579718 | 36774.230941 | 33826.366479 | 18202.822001 | 5306.170778 |
| 168.bin | 31501.506481 | 5333.079725 | 16801.282987 | 26707.248649 | 26168.426756 | 14700.223495 | 4794.257833 |
| 169.bin | 27981.654400 | -233.725069 | 9652.495499 | 20710.786556 | 28215.379469 | 18329.158901 | 7270.867844 |
| 170.bin | 34967.562417 | 1969.548724 | 15198.838098 | 28550.333608 | 32998.013694 | 19768.724319 | 6417.228810 |
| 171.bin | 30230.466688 | 4081.126185 | 15212.003969 | 25233.135827 | 26149.340503 | 15018.462719 | 4997.330861 |
| 172.bin | 35584.848316 | 6979.676920 | 20395.125123 | 31057.161330 | 28605.171397 | 15189.723193 | 4527.686987 |
| 173.bin | 32550.823094 | -829.070440 | 10764.743570 | 24262.446534 | 33379.893534 | 21786.079524 | 8288.376560 |
| 174.bin | 68950.658358 | 33919.932417 | 49359.033233 | 59427.066481 | 35030.725941 | 19591.625125 | 9523.591877 |
| 175.bin | 65024.708106 | 22549.351844 | 40094.216966 | 53792.263158 | 42475.356262 | 24930.491140 | 11232.444948 |
| 176.bin | 68620.320466 | 32915.096848 | 48102.671418 | 57863.498135 | 35705.223617 | 20517.649048 | 10756.822330 |
| 177.bin | 62723.150899 | 27860.149360 | 41725.578599 | 51959.924615 | 34863.001539 | 20997.572300 | 10763.226284 |
| 178.bin | 69178.499088 | 27189.679807 | 43305.466224 | 56880.073913 | 41988.819281 | 25873.032863 | 12298.425174 |
| 179.bin | 64985.865685 | 26498.513224 | 42333.795498 | 54114.583697 | 38487.352461 | 22652.070187 | 10871.281988 |
| 180.bin | 40158.370018 | 9706.994966 | 24802.942101 | 36020.088657 | 30451.375053 | 15355.427917 | 4138.281362 |
| 181.bin | 59905.721557 | 23375.529384 | 36601.152660 | 47885.444661 | 36530.192173 | 23304.568897 | 12020.276896 |
| 182.bin | 60939.242953 | 20913.458809 | 38340.514477 | 51141.151638 | 40025.784144 | 22598.728476 | 9798.091315 |
| 183.bin | 63756.005717 | 27054.090001 | 40418.269366 | 51200.256149 | 36701.915715 | 23337.736351 | 12555.749568 |

| | | | | | | | |
|---|---|---|---|---|---|---|---|
| 184.bin | 26093.287783 | 7128.795861 | 16969.821735 | 23862.669814 | 18964.491921 | 9123.466048 | 2230.617968 |
| 185.bin | 52038.211644 | 12824.493049 | 30171.585687 | 43224.613647 | 39213.718595 | 21866.625957 | 8813.597997 |
| 186.bin | 63659.653988 | 20335.283583 | 38564.112402 | 52284.376481 | 43324.370405 | 25095.541586 | 11375.277507 |
| 187.bin | 68382.537059 | 34389.461557 | 47486.684485 | 57160.706993 | 33993.075501 | 20895.852574 | 11221.830066 |
| 188.bin | 42589.751677 | 8805.766680 | 24925.630912 | 37605.740120 | 33783.984997 | 17664.120766 | 4984.011558 |
| 189.bin | 60356.976807 | 22799.987101 | 39317.761046 | 50631.428256 | 37556.989706 | 21039.215761 | 9725.548551 |
| 18.bin | 59903.714022 | 20822.638336 | 36468.304059 | 48568.978683 | 39081.075686 | 23435.409964 | 11334.735339 |
| 190.bin | 55032.643054 | 22901.765553 | 36809.344834 | 46432.005548 | 32130.877500 | 18223.298220 | 8600.637506 |
| 191.bin | 54971.730505 | 18836.103077 | 34686.299790 | 45807.771423 | 36135.627428 | 20285.430714 | 9163.959082 |
| 192.bin | 45294.625847 | 9388.656672 | 25847.628932 | 38914.905491 | 35905.969175 | 19446.996914 | 6379.720356 |
| 193.bin | 47762.195725 | 9817.414967 | 25691.828740 | 38426.107993 | 37944.780758 | 22070.366985 | 9336.087731 |
| 194.bin | 48909.109407 | 11681.616009 | 26711.716432 | 38708.156509 | 37227.493398 | 22197.392975 | 10200.952898 |
| 195.bin | 57728.176449 | 23144.016307 | 37685.824230 | 48077.875189 | 34584.160142 | 20042.352218 | 9650.301260 |
| 196.bin | 58751.021970 | 18564.081050 | 35535.976131 | 48219.930369 | 40186.940920 | 23215.045839 | 10531.091601 |
| 197.bin | 59062.366183 | 15559.521002 | 34560.448302 | 49412.245050 | 43502.845182 | 24501.917881 | 9650.121133 |
| 198.bin | 59955.015332 | 22701.052730 | 39990.739190 | 50783.097256 | 37253.962602 | 19964.276142 | 9171.918076 |
| 200.bin | 57280.191740 | 23039.687652 | 37425.509416 | 48421.783709 | 34240.504088 | 19854.682324 | 8858.408031 |
| 201.bin | 54772.239392 | 21744.405691 | 36057.398252 | 45780.458192 | 33027.833701 | 18714.841140 | 8991.781200 |
| 202.bin | 55051.735599 | 28837.963992 | 36717.952377 | 44087.019944 | 26213.771608 | 18333.783222 | 10964.715656 |
| 203.bin | 58599.634072 | 18384.696942 | 33834.766566 | 46655.452463 | 40214.937129 | 24764.867506 | 11944.181609 |
| 204.bin | 52605.957143 | 14398.122512 | 31578.131850 | 44659.558665 | 38207.834631 | 21027.825293 | 7946.398478 |
| 205.bin | 62245.464569 | 30999.053565 | 41587.687333 | 50952.224380 | 31246.411004 | 20657.777236 | 11293.240189 |
| 206.bin | 48434.231991 | 11191.931821 | 27709.180220 | 40396.722340 | 37242.300169 | 20725.051771 | 8037.509651 |

| | | | | | | | |
|---|---|---|---|---|---|---|---|
| 207.bin | 58917.845345 | 15317.098263 | 34005.898462 | 48732.961244 | 43600.747082 | 24911.946883 | 10184.884101 |
| 209.bin | 62531.461613 | 23841.483924 | 38650.429146 | 50675.968108 | 38689.977689 | 23881.032467 | 11855.493505 |
| 20.bin | 5345.800724 | -33.968004 | 1800.867540 | 4051.710747 | 5379.768728 | 3544.933184 | 1294.089977 |
| 210.bin | 59405.730500 | 21007.013444 | 37337.985465 | 48731.274422 | 38398.717056 | 22067.745035 | 10674.456078 |
| 211.bin | 84475.055297 | 46970.378001 | 60841.022692 | 71639.442752 | 37504.677296 | 23634.032605 | 12835.612544 |
| 212.bin | 44307.502822 | 8726.201142 | 24766.896921 | 37635.190834 | 35581.301681 | 19540.605901 | 6672.311988 |
| 213.bin | 56753.874231 | 26669.852140 | 37635.513799 | 46212.878145 | 30084.022091 | 19118.360432 | 10540.996086 |
| 214.bin | 59888.883434 | 17741.367163 | 36921.192077 | 50786.667365 | 42147.516271 | 22967.691358 | 9102.216069 |
| 215.bin | 55998.827979 | 16616.842804 | 34687.686154 | 47095.148401 | 39381.985176 | 21311.141825 | 8903.679578 |
| 216.bin | 62718.264988 | 23551.573824 | 38736.719687 | 50750.774907 | 39166.691164 | 23981.545301 | 11967.490081 |
| 218.bin | 6134.469585 | -674.499741 | 1096.536702 | 3466.410993 | 6808.969326 | 5037.932883 | 2668.058592 |
| 22.bin | 36505.655762 | 6145.846986 | 19896.381524 | 31555.675649 | 30359.808775 | 16609.274237 | 4949.980113 |
| 23.bin | 26246.612645 | -904.274631 | 8006.471486 | 18371.656055 | 27150.887276 | 18240.141159 | 7874.956590 |
| 24.bin | 28728.027765 | -6.551929 | 9988.730662 | 21008.775581 | 28734.579695 | 18739.297103 | 7719.252184 |
| 25.bin | 30039.885561 | 588.947991 | 11481.985740 | 23157.370511 | 29450.937570 | 18557.899821 | 6882.515051 |
| 31.bin | 38561.545945 | 5638.415921 | 19683.202129 | 31610.003498 | 32923.130024 | 18878.343816 | 6951.542447 |
| 32.bin | 57293.798605 | 23241.641043 | 37538.089814 | 47416.135972 | 34052.157562 | 19755.708791 | 9877.662633 |
| 33.bin | 56911.226560 | 25624.611219 | 37740.306240 | 46992.973539 | 31286.615341 | 19170.920320 | 9918.253021 |
| 34.bin | 21782.028928 | 2078.269178 | 10033.793206 | 17839.461182 | 19703.759750 | 11748.235721 | 3942.567746 |
| 35.bin | 55307.351030 | 16811.895355 | 34467.068915 | 47117.412303 | 38495.455675 | 20840.282115 | 8189.938728 |
| 38.bin | 351.849620 | 30.716913 | 121.439522 | 221.431938 | 321.132707 | 230.410098 | 130.417682 |
| 39.bin | 57899.974609 | 17569.956689 | 35697.712633 | 49027.856643 | 40330.017919 | 22202.261976 | 8872.117966 |

| | | | | | | | |
|---|---|---|---|---|---|---|---|
| 3.bin | 39150.358273 | 10331.634859 | 24143.965344 | 34474.846514 | 28818.723414 | 15006.392929 | 4675.511760 |
| 40.bin | 39242.109000 | 17277.771895 | 26650.350498 | 32565.008667 | 21964.337105 | 12591.758502 | 6677.100333 |
| 41.bin | 32170.961061 | 5182.854144 | 17433.170394 | 27539.970933 | 26988.106917 | 14737.790667 | 4630.990128 |
| 42.bin | 1869.019562 | 819.478355 | 1277.652995 | 1666.419525 | 1049.541207 | 591.366568 | 202.600037 |
| 44.bin | 637.939241 | 94.269879 | 235.079035 | 475.419331 | 543.669362 | 402.860206 | 162.519910 |
| 45.bin | 114.860238 | -30.911795 | -17.368435 | 32.552103 | 145.772033 | 132.228673 | 82.308135 |
| 46.bin | 58699.711983 | 20065.728351 | 36558.976981 | 49041.685748 | 38633.983632 | 22140.735002 | 9658.026235 |
| 47.bin | 72242.126363 | 39912.369554 | 51429.460420 | 60221.984241 | 32329.756809 | 20812.665943 | 12020.142122 |
| 48.bin | 48397.856326 | 9384.767639 | 26509.204865 | 40505.607550 | 39013.088688 | 21888.651462 | 7892.248777 |
| 4.bin | 56261.065759 | 17931.355231 | 36086.989207 | 48533.205698 | 38329.710528 | 20174.076553 | 7727.860061 |
| 51.bin | 69817.287920 | 34234.140275 | 48981.006294 | 59268.138344 | 35583.147645 | 20836.281627 | 10549.149576 |
| 52.bin | 44999.262782 | 7616.422433 | 24422.518120 | 38033.481950 | 37382.840349 | 20576.744662 | 6965.780832 |
| 54.bin | 50914.623661 | 10607.494201 | 27772.333357 | 41771.250654 | 40307.129460 | 23142.290304 | 9143.373007 |
| 56.bin | 34134.271891 | 4788.149709 | 17203.413777 | 28350.995127 | 29346.122182 | 16930.858114 | 5783.276764 |
| 57.bin | 13746.165878 | 3119.527241 | 7973.977629 | 12111.568208 | 10626.638637 | 5772.188249 | 1634.597670 |
| 58.bin | 62751.822857 | 25539.027091 | 39564.922498 | 50364.352458 | 37212.795766 | 23186.900359 | 12387.470399 |
| 59.bin | 65439.996068 | 24434.602853 | 42163.984313 | 54738.910046 | 41005.393215 | 23276.011755 | 10701.086022 |
| 5.bin | 46122.142258 | 12086.936728 | 27637.329379 | 38807.774404 | 34035.205531 | 18484.812879 | 7314.367854 |
| 60.bin | 60262.476703 | 20032.205653 | 37596.618207 | 50469.221690 | 40230.271049 | 22665.858495 | 9793.255013 |
| 61.bin | 64388.059894 | 28770.256721 | 43159.113285 | 54376.966192 | 35617.803172 | 21228.946608 | 10011.093702 |
| 62.bin | 38115.033863 | 9966.988331 | 24039.089717 | 34551.501938 | 28148.045532 | 14075.944146 | 3563.531925 |
| 63.bin | 29457.417972 | -811.481279 | 9524.202839 | 21621.950282 | 30268.899252 | 19933.215134 | 7835.467691 |
| 64.bin | 59096.927887 | 23020.094705 | 38319.531662 | 48649.427805 | 36076.833182 | 20777.396225 | 10447.500082 |

| | | | | | | | |
|---|---|---|---|---|---|---|---|
| 65.bin | 60256.335848 | 20692.401865 | 37126.169622 | 49449.728234 | 39563.933982 | 23130.166226 | 10806.607614 |
| 66.bin | 494.736878 | -81.107844 | 58.985498 | 285.257187 | 575.844722 | 435.751379 | 209.479690 |
| 67.bin | 58467.954218 | 23827.510433 | 38943.508510 | 48921.058869 | 34640.443785 | 19524.445708 | 9546.895350 |
| 69.bin | 52880.438106 | 8639.332624 | 27624.476428 | 43651.316006 | 44241.105482 | 25255.961678 | 9229.122100 |
| 6.bin | 21490.313801 | 3482.611145 | 11467.275283 | 18562.321994 | 18007.702656 | 10023.038518 | 2927.991807 |
| 71.bin | 52594.320371 | 12693.508566 | 30924.554187 | 44254.609006 | 39900.811805 | 21669.766184 | 8339.711365 |
| 72.bin | 66676.387117 | 29358.532910 | 45700.113828 | 55486.758291 | 37317.854207 | 20976.273289 | 11189.628826 |
| 73.bin | 55126.261428 | 12479.643901 | 31829.880535 | 46556.609115 | 42646.617527 | 23296.380893 | 8569.652313 |
| 74.bin | 23943.100679 | 4315.451699 | 13370.556808 | 21030.484270 | 19627.648980 | 10572.543870 | 2912.616408 |
| 75.bin | 70111.317956 | 33370.947936 | 45265.240214 | 57069.538492 | 36740.370020 | 24846.077742 | 13041.779464 |
| 76.bin | 54496.701659 | 26290.117067 | 38008.561860 | 46142.682810 | 28206.584592 | 16488.139799 | 8354.018849 |
| 77.bin | 61491.984193 | 18180.119071 | 35721.327970 | 49748.416279 | 43311.865123 | 25770.656223 | 11743.567914 |
| 78.bin | 47018.535578 | 15294.040885 | 29353.647493 | 39531.982224 | 31724.494693 | 17664.888085 | 7486.553354 |
| 79.bin | 59419.834523 | 14617.282675 | 35269.217121 | 50376.920186 | 44802.551849 | 24150.617403 | 9042.914337 |
| 7.bin | 55354.010415 | 12544.288304 | 31620.012565 | 46362.487147 | 42809.722111 | 23733.997849 | 8991.523268 |
| 80.bin | 59313.280937 | 17058.637815 | 36262.684564 | 50401.950854 | 42254.643122 | 23050.596374 | 8911.330083 |
| 81.bin | 31341.772509 | -498.791458 | 10839.689681 | 23640.417764 | 31840.563967 | 20502.082828 | 7701.354745 |
| 82.bin | 33694.358416 | 2916.952532 | 15187.172988 | 27060.484362 | 30777.405884 | 18507.185428 | 6633.874054 |
| 83.bin | 31524.506866 | 2462.562746 | 14013.712140 | 25307.653585 | 29061.944120 | 17510.794727 | 6216.853281 |
| 84.bin | 35276.435756 | 7183.609234 | 20232.161792 | 30523.361488 | 28092.826523 | 15044.273964 | 4753.074268 |
| 85.bin | 62783.078353 | 19702.217017 | 37552.596713 | 50538.631618 | 43080.861336 | 25230.481639 | 12244.446735 |
| 86.bin | 21178.923141 | 4072.488610 | 12036.348534 | 18686.066326 | 17106.434531 | 9142.574607 | 2492.856815 |
| 87.bin | 60670.934221 | 30358.637416 | 39836.195325 | 48997.566125 | 30312.296805 | 20834.738896 | 11673.368096 |

| | | | | | | | |
|---|---|---|---|---|---|---|---|
| 88.bin | 76145.318000 | 40848.634955 | 54055.761327 | 64210.820894 | 35296.683045 | 22089.556673 | 11934.497106 |
| 89.bin | 63635.382021 | 29595.709209 | 41788.311935 | 52631.985630 | 34039.672812 | 21847.070086 | 11003.396391 |
| 8.bin | 60460.938347 | 28098.639585 | 39431.022474 | 49279.743313 | 32362.298763 | 21029.915873 | 11181.195034 |
| 90.bin | 59316.475753 | 18368.634653 | 35055.708284 | 48277.452622 | 40947.841100 | 24260.767468 | 11039.023131 |
| 91.bin | 61764.313226 | 31069.138061 | 40253.442863 | 50409.412129 | 30695.175165 | 21510.870363 | 11354.901097 |
| 92.bin | 45668.880578 | 7583.384259 | 24872.620884 | 38852.995933 | 38085.496319 | 20796.259694 | 6815.884645 |
| 93.bin | 60722.777693 | 26982.185315 | 40973.902088 | 51195.660208 | 33740.592377 | 19748.875605 | 9527.117485 |
| 94.bin | 5321.276106 | 819.699187 | 2392.042551 | 4071.874613 | 4501.576919 | 2929.233555 | 1249.401493 |
| 95.bin | 15677.854753 | 2608.977694 | 7754.909776 | 12407.129672 | 13068.877059 | 7922.944977 | 3270.725081 |
| 97.bin | 547.951181 | 42.062997 | 146.828366 | 289.239995 | 505.888184 | 401.122815 | 258.711187 |
| 98.bin | 2300.252461 | 163.731459 | 939.050966 | 1810.745541 | 2136.521002 | 1361.201496 | 489.506920 |
| 99.bin | 42520.984656 | 11056.037604 | 25642.909802 | 36134.412759 | 31464.947052 | 16878.074855 | 6386.571897 |
| 9.bin | 76453.981701 | 41515.695247 | 51307.147349 | 63774.155500 | 34938.286454 | 25146.834352 | 12679.826201 |

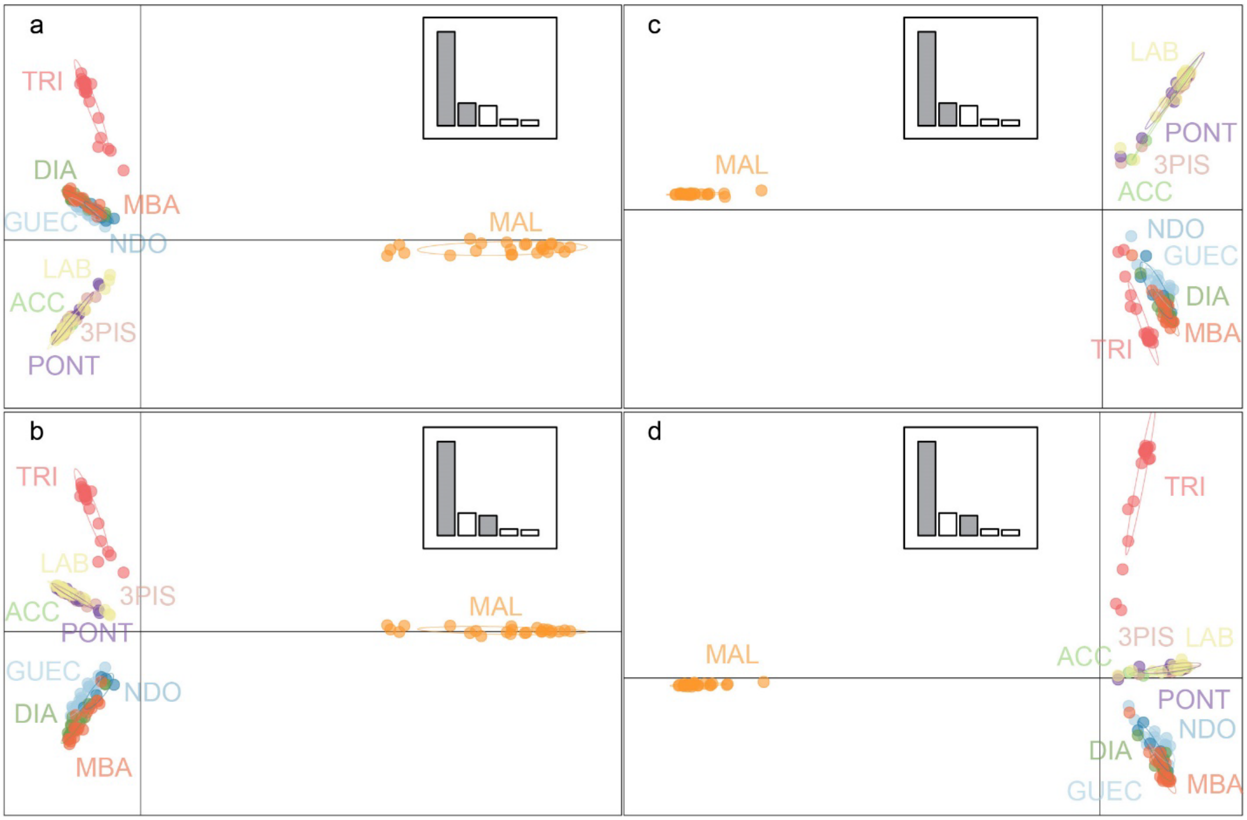


**Figure A1 The PCA analyses of the diploid (a, b) and tetraploid (c,d) datasets.** a): PCA axes 1 and 2 and b) PCA axes 1 and 3 of the diploid dataset; c) PCA axes 1 and 2 and d) PCA axes 1 and 3 for the tetraploid dataset. Different colours indicate different sampling sites. The full names of the abbreviations can be found in Table 1.

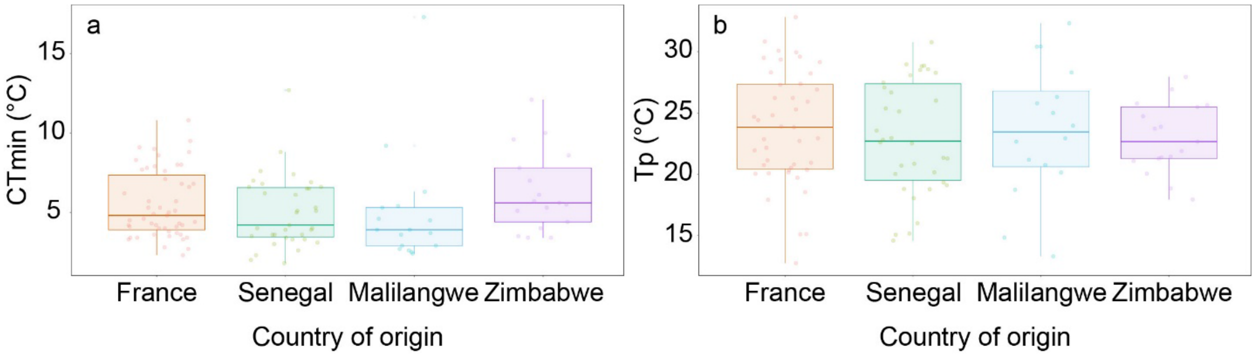


**Figure A2 Results of the life history analyses**. a) $CT_{min}$ and b) $T_p$ as a function of country of origin. No significant effects of country of origin could be detected.

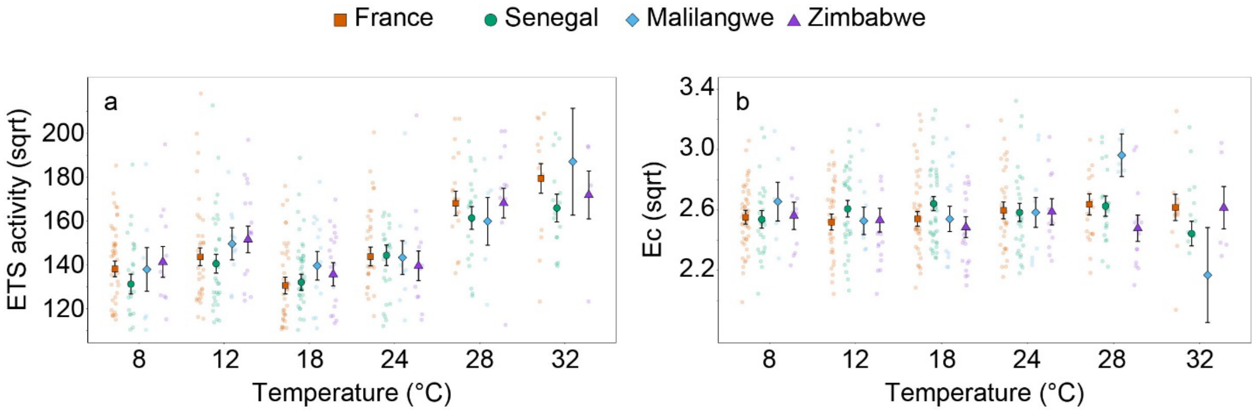


**Figure A3 Results of the physiological analyses**. a) ETS activity in nmol O2/min/mg protein and b) the total energy consumption as a function of temperature and country of origin. No significant effects of country of origin could be detected.

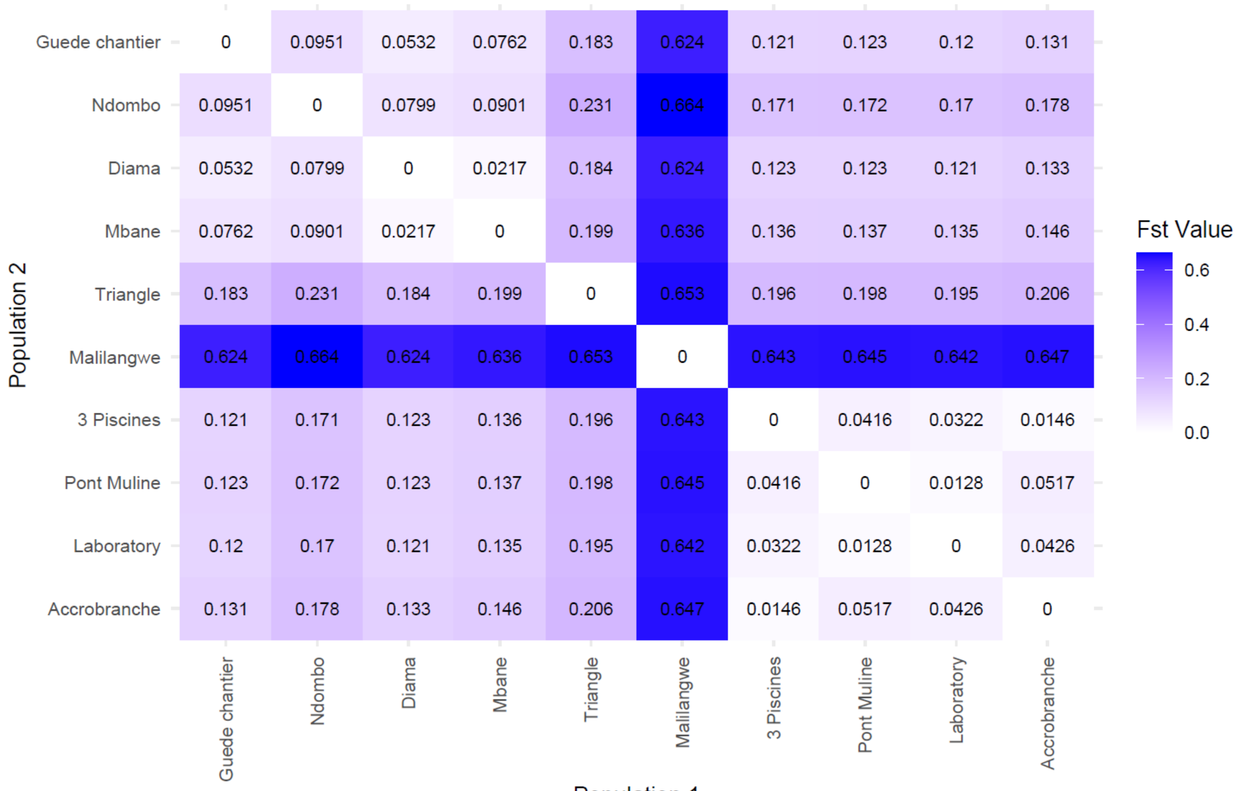


**Figure A4 Pairwise $F_{ST}$ values between the different populations for the neutral dataset.**

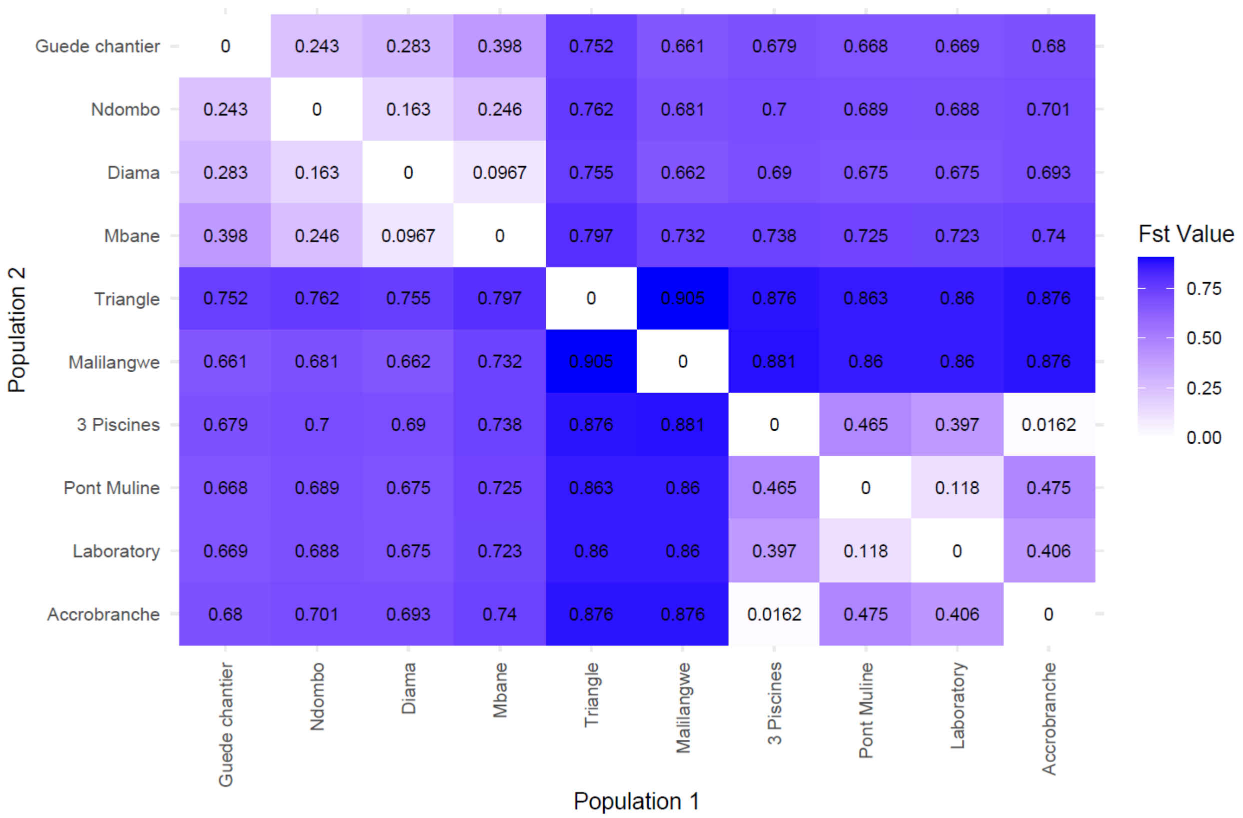


| | Guede chantier | Ndombo | Diama | Mbane | Triangle | Malilangwe | 3 Piscines | Pont Muline | Laboratory | Accrobranche |
|---|---|---|---|---|---|---|---|---|---|---|
| Guede chantier | 0 | 0.243 | 0.283 | 0.398 | 0.752 | 0.661 | 0.679 | 0.668 | 0.669 | 0.68 |
| Ndombo | 0.243 | 0 | 0.163 | 0.246 | 0.762 | 0.681 | 0.7 | 0.689 | 0.688 | 0.701 |
| Diama | 0.283 | 0.163 | 0 | 0.0967 | 0.755 | 0.662 | 0.69 | 0.675 | 0.675 | 0.693 |
| Mbane | 0.398 | 0.246 | 0.0967 | 0 | 0.797 | 0.732 | 0.738 | 0.725 | 0.723 | 0.74 |
| Triangle | 0.752 | 0.762 | 0.755 | 0.797 | 0 | 0.905 | 0.876 | 0.863 | 0.86 | 0.876 |
| Malilangwe | 0.661 | 0.681 | 0.662 | 0.732 | 0.905 | 0 | 0.881 | 0.86 | 0.86 | 0.876 |
| 3 Piscines | 0.679 | 0.7 | 0.69 | 0.738 | 0.876 | 0.881 | 0 | 0.465 | 0.397 | 0.0162 |
| Pont Muline | 0.668 | 0.689 | 0.675 | 0.725 | 0.863 | 0.86 | 0.465 | 0 | 0.118 | 0.475 |
| Laboratory | 0.669 | 0.688 | 0.675 | 0.723 | 0.86 | 0.86 | 0.397 | 0.118 | 0 | 0.406 |
| Accrobranche | 0.68 | 0.701 | 0.693 | 0.74 | 0.876 | 0.876 | 0.0162 | 0.475 | 0.406 | 0 |

**Figure A5 Pairwise $F_{ST}$ values between the different populations for the outlier dataset.**

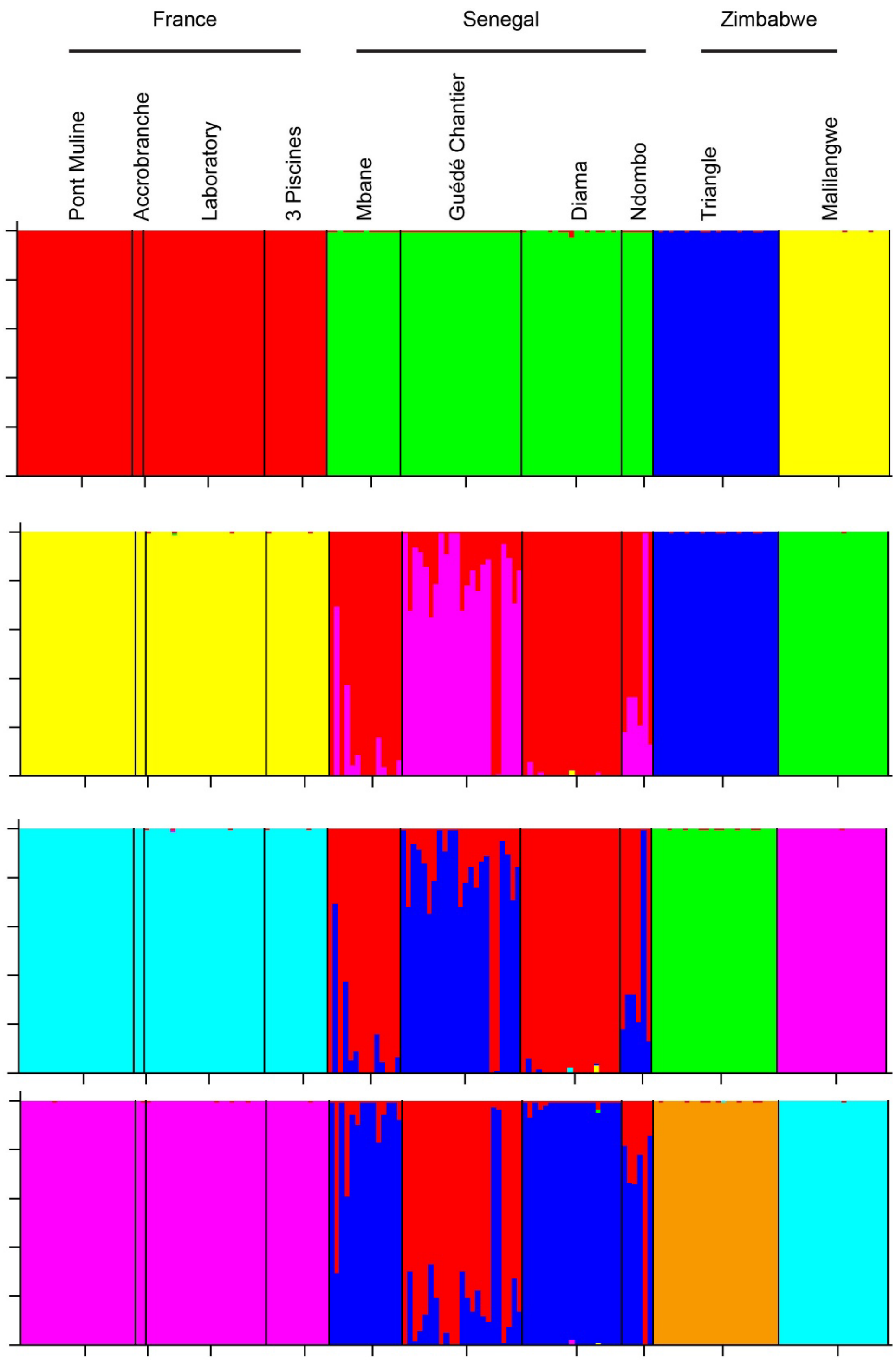


**Figure A6 Results of the STRUCTURE analyses for a) K=4, b) K=5, c) K=6 and d) K=7.** The optimal number of populations was K=5, thereby following the method of Evanno et al. (2005). The x-axis shows the different populations and the corresponding population is given on the right.

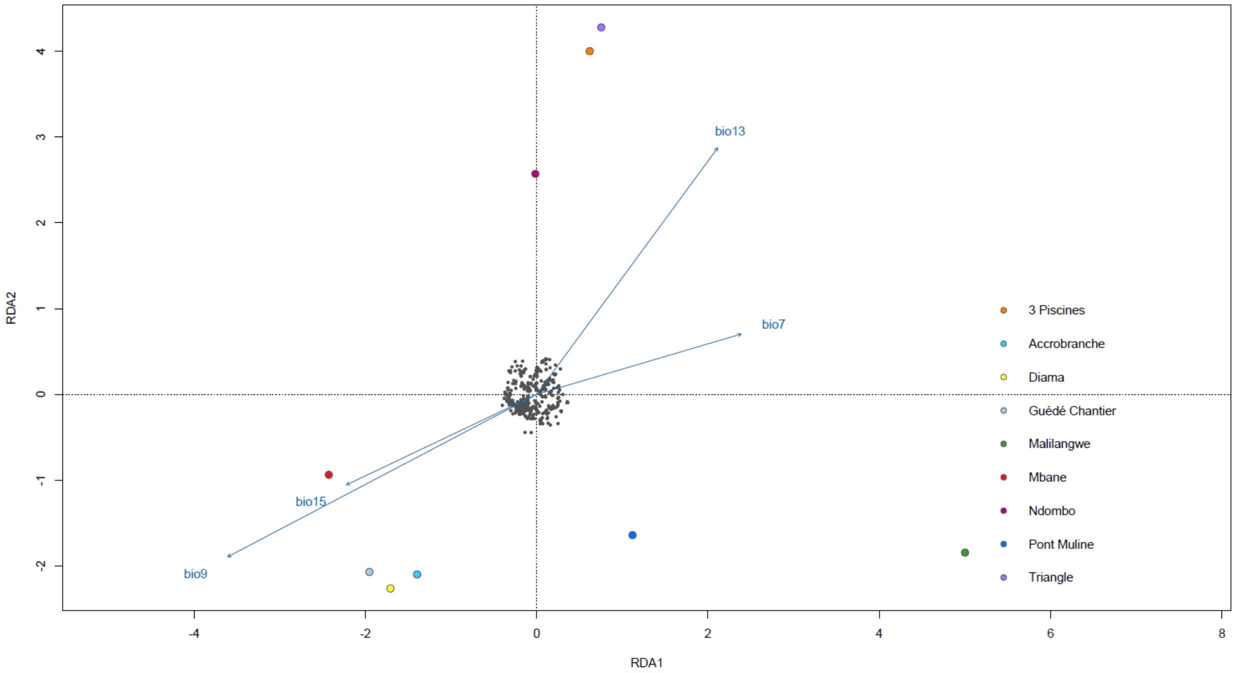


**Figure A7 RDA analysis with the climatic data on the outlier dataset.** bio7 and bio 9 are temperature-related variables, bio 13 and bio 15 are precipitation-related variables.

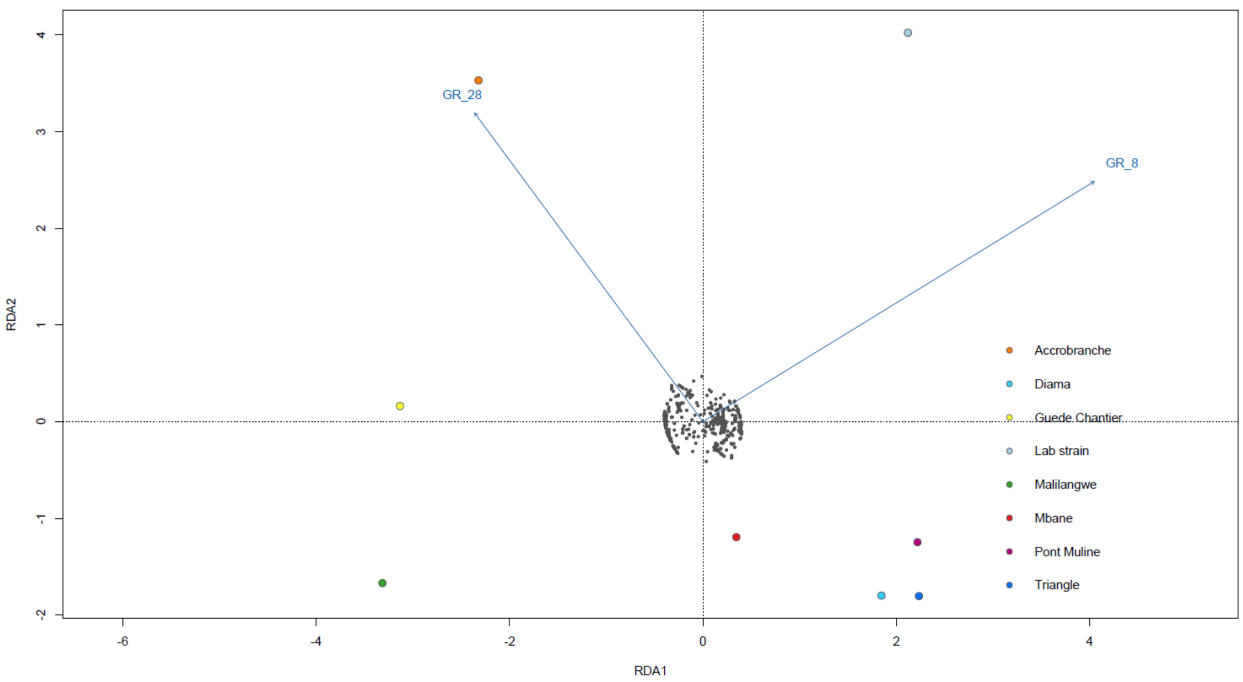


**Figure A8 RDA analysis with the life history data on the outlier dataset.** growth rate at 8 °C (GR_8) and growth rate at 28 °C (GR_28) are most likely associated with some of the outlier SNPs.

**Table A1 The number of snails per locality and temperature treatment included for analysis of each physiological variable**

| Locality | Country | Temperature | µg fat/mg sample | µg protein /mg sample | µg sugar /mg sample | O2/mg sample | Nmol MDA /mg fat | Hemoglobin/mg protein | Nmol PO/min/ mg protein | CEA |
|---|---|---|---|---|---|---|---|---|---|---|
| **Diama** | Senegal | 4 | 0 | 0 | 0 | 0 | 0 | 0 | 0 | 0 |
| **Mbane** | Senegal | 4 | 0 | 0 | 0 | 0 | 0 | 0 | 0 | 0 |
| **Guede-Chantier** | Senegal | 4 | 0 | 0 | 0 | 0 | 0 | 0 | 0 | 0 |
| **Triangle** | Zimbabwe | 4 | 0 | 0 | 0 | 0 | 0 | 0 | 0 | 0 |
| **Malilangwe** | Zimbabwe | 4 | 0 | 0 | 0 | 0 | 0 | 0 | 0 | 0 |
| **Pont Muline** | Corsica | 4 | 4 | 4 | 4 | 4 | 4 | 4 | 4 | 4 |
| **Accrobranche** | Corsica | 4 | 0 | 0 | 0 | 0 | 0 | 0 | 0 | 0 |
| **Lab strain** | Corsica | 4 | 0 | 0 | 0 | 0 | 0 | 0 | 0 | 0 |
| **Diama** | Senegal | 8 | 13 | 13 | 13 | 13 | 13 | 13 | 13 | 13 |
| **Mbane** | Senegal | 8 | 8 | 8 | 8 | 8 | 8 | 8 | 8 | 8 |
| **Guede-Chantier** | Senegal | 8 | 8 | 8 | 8 | 8 | 8 | 8 | 8 | 8 |
| **Triangle** | Zimbabwe | 8 | 12 | 12 | 12 | 12 | 12 | 12 | 11 | 12 |
| **Malilangwe** | Zimbabwe | 8 | 6 | 6 | 6 | 6 | 6 | 6 | 6 | 6 |
| **Pont Muline** | Corsica | 8 | 22 | 22 | 22 | 22 | 21 | 22 | 19 | 22 |
| **Accrobranche** | Corsica | 8 | 3 | 3 | 3 | 3 | 3 | 3 | 3 | 3 |
| **Lab strain** | Corsica | 8 | 21 | 22 | 20 | 22 | 21 | 22 | 18 | 20 |
| **Diama** | Senegal | 12 | 11 | 11 | 11 | 11 | 11 | 11 | 11 | 11 |
| **Mbane** | Senegal | 12 | 12 | 12 | 12 | 12 | 12 | 12 | 11 | 12 |
| **Guede-Chantier** | Senegal | 12 | 11 | 11 | 11 | 10 | 11 | 11 | 11 | 10 |
| **Triangle** | Zimbabwe | 12 | 16 | 16 | 14 | 16 | 16 | 16 | 15 | 14 |
| **Malilangwe** | Zimbabwe | 12 | 12 | 11 | 12 | 11 | 12 | 11 | 11 | 11 |
| **Pont Muline** | Corsica | 12 | 19 | 19 | 19 | 19 | 19 | 19 | 18 | 19 |
| **Accrobranche** | Corsica | 12 | 5 | 5 | 5 | 5 | 5 | 5 | 5 | 5 |
| **Lab strain** | Corsica | 12 | 12 | 12 | 12 | 12 | 12 | 12 | 12 | 12 |
| **Diama** | Senegal | 18 | 18 | 18 | 18 | 18 | 18 | 18 | 18 | 18 |
| **Mbane** | Senegal | 18 | 14 | 14 | 14 | 14 | 14 | 14 | 14 | 14 |

| | | | | | | | | | | |
|---|---|---|---|---|---|---|---|---|---|---|
| **Guede-Chantier** | Senegal | 18 | 13 | 13 | 13 | 13 | 12 | 13 | 13 | 13 |
| **Triangle** | Zimbabwe | 18 | 21 | 21 | 21 | 21 | 21 | 21 | 21 | 21 |
| **Malilangwe** | Zimbabwe | 18 | 14 | 14 | 14 | 14 | 14 | 14 | 13 | 14 |
| **Pont Muline** | Corsica | 18 | 20 | 20 | 20 | 20 | 20 | 20 | 20 | 20 |
| **Accrobranche** | Corsica | 18 | 6 | 6 | 6 | 6 | 6 | 6 | 6 | 6 |
| **Lab strain** | Corsica | 18 | 16 | 15 | 16 | 15 | 16 | 15 | 15 | 15 |
| **Diama** | Senegal | 24 | 8 | 8 | 8 | 8 | 8 | 8 | 8 | 8 |
| **Mbane** | Senegal | 24 | 9 | 9 | 9 | 9 | 9 | 9 | 9 | 9 |
| **Guede-Chantier** | Senegal | 24 | 10 | 10 | 10 | 10 | 10 | 10 | 10 | 10 |
| **Triangle** | Zimbabwe | 24 | 13 | 13 | 13 | 13 | 13 | 13 | 13 | 13 |
| **Malilangwe** | Zimbabwe | 24 | 10 | 10 | 10 | 10 | 10 | 10 | 10 | 10 |
| **Pont Muline** | Corsica | 24 | 15 | 15 | 15 | 15 | 15 | 15 | 15 | 15 |
| **Accrobranche** | Corsica | 24 | 3 | 3 | 3 | 3 | 3 | 3 | 3 | 3 |
| **Lab strain** | Corsica | 24 | 14 | 14 | 14 | 14 | 14 | 14 | 14 | 14 |
| **Diama** | Senegal | 28 | 8 | 8 | 8 | 8 | 8 | 8 | 8 | 8 |
| **Mbane** | Senegal | 28 | 9 | 9 | 9 | 9 | 9 | 9 | 9 | 9 |
| **Guede-Chantier** | Senegal | 28 | 5 | 5 | 5 | 5 | 5 | 5 | 5 | 5 |
| **Triangle** | Zimbabwe | 28 | 13 | 13 | 13 | 13 | 13 | 13 | 13 | 13 |
| **Malilangwe** | Zimbabwe | 28 | 5 | 5 | 5 | 5 | 5 | 5 | 5 | 5 |
| **Pont Muline** | Corsica | 28 | 10 | 10 | 10 | 10 | 10 | 10 | 10 | 10 |
| **Accrobranche** | Corsica | 28 | 2 | 2 | 2 | 2 | 2 | 2 | 2 | 2 |
| **Lab strain** | Corsica | 28 | 8 | 8 | 8 | 8 | 8 | 8 | 7 | 8 |
| **Diama** | Senegal | 32 | 5 | 6 | 6 | 6 | 5 | 6 | 6 | 5 |
| **Mbane** | Senegal | 32 | 6 | 6 | 6 | 6 | 6 | 6 | 6 | 6 |
| **Guede-Chantier** | Senegal | 32 | 3 | 3 | 3 | 3 | 3 | 3 | 3 | 3 |
| **Triangle** | Zimbabwe | 32 | 5 | 5 | 5 | 5 | 5 | 5 | 5 | 5 |
| **Malilangwe** | Zimbabwe | 32 | 1 | 1 | 1 | 1 | 1 | 1 | 1 | 1 |
| **Pont Muline** | Corsica | 32 | 1 | 1 | 1 | 1 | 1 | 1 | 1 | 1 |
| **Accrobranche** | Corsica | 32 | 1 | 1 | 1 | 1 | 1 | 1 | 1 | 1 |

| | | | | | | | | | | |
|---|---|---|---|---|---|---|---|---|---|---|
| **Lab strain** | Corsica | 32 | 8 | 8 | 8 | 8 | 8 | 8 | 7 | 8 |

**Table A2 Basic population statistics for each population that was included in the genetic analysis for both the neutral and the outlier dataset.** Ar stands for allelic richness, Ho for observed heterozygosity, He for expected heterozygosity, and Fis for inbreeding coefficient

| | Neutral dataset | | | | Outlier dataset | | | |
|---|---|---|---|---|---|---|---|---|
| **Population** | Ar | Ho | He | Fis | Ar | Ho | He | Fis |
| **Guédé Chantier** | 1.368 | 0.618 | 0.360 | -0.659 | 1.190 | 0.197 | 0.190 | 0.0744 |
| **Ndombo** | 1.386 | 0.473 | 0.356 | -0.349 | 1.225 | 0.201 | 0.232 | 0.0527 |
| **Diama** | 1.374 | 0.628 | 0.361 | -0.678 | 1.199 | 0.227 | 0.197 | -0.0502 |
| **Triangle** | 1.345 | 0.608 | 0.336 | -0.796 | 1.050 | 0.0859 | 0.0486 | -0.572 |
| **Mbane** | 1.358 | 0.592 | 0.349 | -0.643 | 1.159 | 0.214 | 0.156 | -0.230 |
| **Malilangwe** | 1.006 | 0.00977 | 0.006133 | -0.243 | 1.00009 | 9.1e-05 | 9.1e-05 | 0 |
| **3 Piscines** | 1.346 | 0.642 | 0.337 | -0.893 | 1.0670 | 0.124 | 0.0653 | -0.882 |
| **Accrobranche** | 1.501 | 0.650 | 0.346 | -0.913 | 1.099 | 0.126 | 0.0606 | -0.923 |
| **Pont Mulinu** | 1.355 | 0.615 | 0.339 | -0.794 | 1.0858 | 0.127 | 0.0835 | -0.326 |
| **Laboratory** | 1.347 | 0.616 | 0.339 | -0.797 | 1.0863 | 0.128 | 0.0850 | -0.383 |

**Table A3 loci associated with the different bioclimatic variables.** bio7 is the temperature annual range, bio9 is the mean temperature of the driest quarter, bio13 is the precipitation of the wettest month, and bio15 is the precipitation seasonality.

| Locus | Associated with variables | | |
|---|---|---|---|
| JAGDYQ010000001_1_31892982 | bio9 | bio15 | |
| JAGDYQ010000002_1_14338404 | bio15 | | |
| JAGDYQ010000002_1_4670299 | bio15 | | |
| JAGDYQ010000004_1_12391587 | bio9 | | |
| JAGDYQ010000004_1_4894076 | bio15 | | |
| JAGDYQ010000004_1_4894081 | bio9 | | |
| JAGDYQ010000004_1_4894443 | bio9 | | |
| JAGDYQ010000004_1_4895125 | bio15 | | |
| JAGDYQ010000004_1_5499892 | bio15 | | |
| JAGDYQ010000004_1_6252191 | bio9 | bio13 | bio15 |
| JAGDYQ010000005_1_5080124 | bio15 | | |
| JAGDYQ010000006_1_12391320 | bio7 | bio9 | bio15 |
| JAGDYQ010000007_1_11486841 | bio15 | | |
| JAGDYQ010000007_1_4737766 | bio9 | | |
| JAGDYQ010000007_1_6590701 | bio15 | | |
| JAGDYQ010000007_1_7904848 | bio15 | | |
| JAGDYQ010000008_1_10958326 | bio9 | | |
| JAGDYQ010000008_1_12717101 | bio15 | | |
| JAGDYQ010000010_1_6442779 | bio13 | bio15 | |
| JAGDYQ010000010_1_7648562 | bio9 | bio15 | |
| JAGDYQ010000012_1_2928984 | bio9 | bio13 | bio15 |
| JAGDYQ010000012_1_4355478 | bio9 | | |
| JAGDYQ010000012_1_4406528 | bio9 | | |
| JAGDYQ010000014_1_7750482 | bio9 | | |
| JAGDYQ010000016_1_2919255 | bio9 | | |
| JAGDYQ010000017_1_2039233 | bio15 | | |
| JAGDYQ010000018_1_8553059 | bio15 | | |
| JAGDYQ010000018_1_9508614 | bio9 | | |
| JAGDYQ010000019_1_3011795 | bio15 | | |
| JAGDYQ010000019_1_4112529 | bio15 | | |
| JAGDYQ010000019_1_9286789 | bio15 | | |
| JAGDYQ010000021_1_96115 | bio15 | | |
| JAGDYQ010000023_1_5093997 | bio15 | | |
| JAGDYQ010000024_1_7689858 | bio9 | | |
| JAGDYQ010000024_1_8638883 | bio9 | | |
| JAGDYQ010000024_1_8638911 | bio15 | | |
| JAGDYQ010000026_1_4553200 | bio9 | | |
| JAGDYQ010000028_1_4429165 | bio15 | | |
| JAGDYQ010000029_1_6428458 | bio9 | | |
| JAGDYQ010000029_1_7030517 | bio9 | | |
| JAGDYQ010000029_1_8419841 | bio15 | | |
| JAGDYQ010000030_1_4062784 | bio15 | | |
| JAGDYQ010000030_1_4434253 | bio9 | | |

| | | | |
|---|---|---|---|
| JAGDYQ010000033_1_2404233 | bio15 | | |
| JAGDYQ010000033_1_3926816 | bio15 | | |
| JAGDYQ010000036_1_2746453 | bio9 | | |
| JAGDYQ010000038_1_4275692 | bio15 | | |
| JAGDYQ010000038_1_6560426 | bio15 | | |
| JAGDYQ010000044_1_3732115 | bio15 | | |
| JAGDYQ010000045_1_3358886 | bio9 | bio13 | bio15 |
| JAGDYQ010000047_1_5830196 | bio7 | bio9 | |
| JAGDYQ010000047_1_5830212 | bio7 | bio9 | |
| JAGDYQ010000047_1_5830216 | bio7 | bio9 | |
| JAGDYQ010000053_1_2597612 | bio15 | | |
| JAGDYQ010000054_1_2833202 | bio9 | | |
| JAGDYQ010000054_1_3570088 | bio9 | | |
| JAGDYQ010000059_1_1992894 | bio9 | | |
| JAGDYQ010000059_1_306443 | bio9 | | |
| JAGDYQ010000059_1_3106777 | bio9 | | |
| JAGDYQ010000059_1_922013 | bio9 | bio15 | |
| JAGDYQ010000060_1_5060434 | bio9 | | |
| JAGDYQ010000060_1_5060444 | bio9 | | |
| JAGDYQ010000062_1_4224762 | bio9 | | |
| JAGDYQ010000064_1_186743 | bio15 | | |
| JAGDYQ010000065_1_2842779 | bio15 | | |
| JAGDYQ010000069_1_824600 | bio7 | bio9 | |
| JAGDYQ010000072_1_1217840 | bio15 | | |
| JAGDYQ010000078_1_1534621 | bio15 | | |
| JAGDYQ010000080_1_198295 | bio15 | | |
| JAGDYQ010000080_1_709862 | bio9 | bio15 | |
| JAGDYQ010000081_1_3011198 | bio15 | | |
| JAGDYQ010000081_1_4248240 | bio9 | | |
| JAGDYQ010000082_1_3090146 | bio9 | | |
| JAGDYQ010000083_1_1041457 | bio9 | bio15 | |
| JAGDYQ010000083_1_2648714 | bio15 | | |
| JAGDYQ010000085_1_2709680 | bio9 | | |
| JAGDYQ010000086_1_354296 | bio9 | | |
| JAGDYQ010000086_1_354307 | bio9 | | |
| JAGDYQ010000086_1_354338 | bio9 | | |
| JAGDYQ010000086_1_354339 | bio13 | bio15 | |
| JAGDYQ010000086_1_406933 | bio15 | | |
| JAGDYQ010000090_1_751964 | bio9 | | |
| JAGDYQ010000090_1_751979 | bio9 | | |
| JAGDYQ010000091_1_2216813 | bio9 | | |
| JAGDYQ010000091_1_3999503 | bio9 | bio15 | |
| JAGDYQ010000092_1_2772241 | bio9 | | |
| JAGDYQ010000093_1_153601 | bio9 | | |
| JAGDYQ010000093_1_2875635 | bio15 | | |
| JAGDYQ010000098_1_3975170 | bio9 | | |
| JAGDYQ010000100_1_8069 | bio9 | | |
| JAGDYQ010000101_1_1679108 | bio15 | | |

| | | | | |
|---|---|---|---|---|
| JAGDYQ010000101_1_1730119 | bio9 | | | |
| JAGDYQ010000101_1_1939810 | bio9 | | | |
| JAGDYQ010000105_1_965900 | bio15 | | | |
| JAGDYQ010000108_1_1407366 | bio15 | | | |
| JAGDYQ010000108_1_2110576 | bio13 | | | |
| JAGDYQ010000110_1_1017499 | bio15 | | | |
| JAGDYQ010000110_1_1017532 | bio15 | | | |
| JAGDYQ010000110_1_614001 | bio9 | bio15 | | |
| JAGDYQ010000113_1_1407145 | bio9 | | | |
| JAGDYQ010000119_1_2932088 | bio9 | | | |
| JAGDYQ010000120_1_1671578 | bio9 | | | |
| JAGDYQ010000120_1_1671583 | bio9 | | | |
| JAGDYQ010000122_1_2955356 | bio15 | | | |
| JAGDYQ010000123_1_206057 | bio15 | | | |
| JAGDYQ010000128_1_2170263 | bio15 | | | |
| JAGDYQ010000128_1_2787144 | bio9 | | | |
| JAGDYQ010000129_1_1637206 | bio15 | | | |
| JAGDYQ010000129_1_2867165 | bio15 | | | |
| JAGDYQ010000132_1_1815190 | bio15 | | | |
| JAGDYQ010000134_1_482688 | bio9 | bio13 | bio15 | |
| JAGDYQ010000134_1_483117 | bio9 | | | |
| JAGDYQ010000136_1_377912 | bio15 | | | |
| JAGDYQ010000137_1_432115 | bio9 | | | |
| JAGDYQ010000141_1_1052893 | bio9 | | | |
| JAGDYQ010000142_1_971187 | bio15 | | | |
| JAGDYQ010000143_1_1405194 | bio9 | | | |
| JAGDYQ010000151_1_2457706 | bio15 | | | |
| JAGDYQ010000151_1_901764 | bio15 | | | |
| JAGDYQ010000156_1_1023583 | bio9 | | | |
| JAGDYQ010000156_1_1023602 | bio9 | | | |
| JAGDYQ010000159_1_2331879 | bio9 | bio15 | | |
| JAGDYQ010000161_1_1459792 | bio9 | | | |
| JAGDYQ010000161_1_1459817 | bio9 | bio15 | | |
| JAGDYQ010000161_1_594978 | bio9 | | | |
| JAGDYQ010000162_1_1264298 | bio15 | | | |
| JAGDYQ010000162_1_29342 | bio7 | bio9 | bio13 | bio15 |
| JAGDYQ010000165_1_495165 | bio9 | | | |
| JAGDYQ010000178_1_1700830 | bio15 | | | |
| JAGDYQ010000178_1_187242 | bio15 | | | |
| JAGDYQ010000183_1_1750880 | bio15 | | | |
| JAGDYQ010000202_1_1521865 | bio9 | bio15 | | |
| JAGDYQ010000206_1_1351171 | bio15 | | | |
| JAGDYQ010000218_1_1232108 | bio9 | | | |
| JAGDYQ010000222_1_1391428 | bio9 | | | |
| JAGDYQ010000234_1_1091160 | bio9 | | | |
| JAGDYQ010000235_1_1196080 | bio9 | | | |
| JAGDYQ010000250_1_175453 | bio9 | bio15 | | |
| JAGDYQ010000256_1_836697 | bio9 | | | |

| | | | |
|---|---|---|---|
| JAGDYQ010000258_1_1012643 | bio9 | | |
| JAGDYQ010000267_1_79449 | bio15 | | |
| JAGDYQ010000267_1_79450 | bio15 | | |
| JAGDYQ010000267_1_79451 | bio9 | bio15 | |
| JAGDYQ010000268_1_924597 | bio15 | | |
| JAGDYQ010000277_1_497056 | bio9 | | |
| JAGDYQ010000277_1_681887 | bio15 | | |
| JAGDYQ010000286_1_188629 | bio9 | bio13 | bio15 |
| JAGDYQ010000296_1_330820 | bio15 | | |
| JAGDYQ010000304_1_544530 | bio9 | | |
| JAGDYQ010000325_1_439481 | bio9 | | |
| JAGDYQ010000331_1_433751 | bio9 | | |
| JAGDYQ010000337_1_24794 | bio9 | | |
| JAGDYQ010000345_1_15934 | bio15 | | |
| JAGDYQ010000351_1_354195 | bio9 | | |
| JAGDYQ010000358_1_180180 | bio15 | | |
| JAGDYQ010000367_1_471806 | bio9 | bio15 | |
| JAGDYQ010000373_1_368929 | bio7 | bio9 | |
| JAGDYQ010000374_1_433574 | bio9 | | |
| JAGDYQ010000374_1_433593 | bio9 | | |
| JAGDYQ010000376_1_175517 | bio7 | bio9 | bio15 |
| JAGDYQ010000388_1_398608 | bio13 | | |
| JAGDYQ010000406_1_108401 | bio15 | | |
| JAGDYQ010000431_1_222650 | bio9 | | |
| JAGDYQ010000432_1_15803 | bio9 | | |
| JAGDYQ010000434_1_213726 | bio9 | | |
